# A neuronal circuit for vector computation builds an allocentric traveling-direction signal in the *Drosophila* fan-shaped body

**DOI:** 10.1101/2020.12.22.423967

**Authors:** Cheng Lyu, L.F. Abbott, Gaby Maimon

## Abstract

Many behavioral tasks require the manipulation of mathematical vectors, but, outside of computational models^1–8^, it is not known how brains perform vector operations. Here we show how the *Drosophila* central complex, a region implicated in goal-directed navigation^8–14^, performs vector arithmetic. First, we describe neural signals in the fan-shaped body that explicitly track a fly’s *allocentric* traveling direction, that is, the traveling direction in reference to external cues. Past work has identified neurons in *Drosophila*^12,15–17^ and mammals^18,19^ that track allocentric heading (e.g., head-direction cells), but these new signals illuminate how the sense of space is properly updated when traveling and heading angles differ. We then characterize a neuronal circuit that rotates, scales, and adds four vectors related to the fly’s *egocentric* traveling direction–– the traveling angle referenced to the body axis––to compute the allocentric traveling direction. Each two-dimensional vector is explicitly represented by a sinusoidal activity pattern across a distinct neuronal population, with the sinusoid’s amplitude representing the vector’s length and its phase representing the vector’s angle. The principles of this circuit, which performs an egocentric-to-allocentric coordinate transformation, may generalize to other brains and to domains beyond navigation where vector operations or reference-frame transformations are required.

## Introduction

Navigating animals often construct an internal sense of their position and orientation in an environment^20^. In mammals, cellular activity correlates of this internal sense include the responses of head direction cells^18^, place cells^21^, grid cells^22^, speed cells^23^, and others. Understanding how such neural signals are built and how they impact behavior is a central goal of cognitive neuroscience.

A fundamental open question in navigation is how an animal computes its direction of travel when that direction is not aligned with its head or body^24–27^. This issue is particularly acute when animals fly or swim because flow forces (e.g. wind) can induce significant sideward and even backward drift. Misalignment of the heading and traveling directions is also relevant for terrestrial locomotion when an animal walks sideways or backwards^27^. Head-direction cells signal the angle the head is facing relative to external landmarks, not the direction the body is moving referenced to those landmarks. Traveling-direction tuned neurons, should they exist, would provide a more generally useful signal for navigation.

Insects solve remarkable navigational tasks^28–30^ and, like mammals, they have head-direction-like cells called ***EPGs***, *compass neurons* or *heading neurons* with activity tuned to the animal’s head (or body) angle in reference to external cues^9,12,31,32^. In *Drosophila*, sixteen sets of EPGs tile the circular ellipsoid body and generate, as a population, a bolus or bump of activity at a position around the ellipsoid body that reflects the fly’s orientation, akin to a compass needle^12,33^. Here we show that two populations of neurons that tile the fan-shaped body––another prominent central-complex structure––each express a bolus of activity at a position along this structure that indicates the fly’s traveling angle in reference to external cues, rather than its heading angle. We experimentally define and theoretically analyze a neural circuit that computes this external-cue-referenced, or *allocentric*, traveling-direction signal by rotating, scaling and summing four vectors related to the fly’s body-axis-referenced, or *egocentric*, traveling direction. Each vector is explicitly represented by a sinusoidal pattern of activity across a defined population of neurons. Characterizing a neuronal circuit for vector arithmetic and reference-frame transformation is a fundamental advance for cognitive neuroscience^34–36^.

### Beyond heading signals in the central complex

In the *Drosophila* central complex (Fig. 1a), single EPG neurons receive synaptic input in one of sixteen wedges that form the ellipsoid body, and they send axonal outputs to one of eighteen glomeruli that form the protocerebral bridge (Fig. 1b, blue cells). In both walking^12,15,16^ and flying flies^14,37^, the full population of EPG cells expresses a single localized bolus of activity in the ellipsoid body, alongside one copy of this bolus in the left protocerebral bridge and another copy in the right protocerebral bridge, at positions along these structures that signal the fly’s angular heading referenced to external cues.

**Figure 1.**
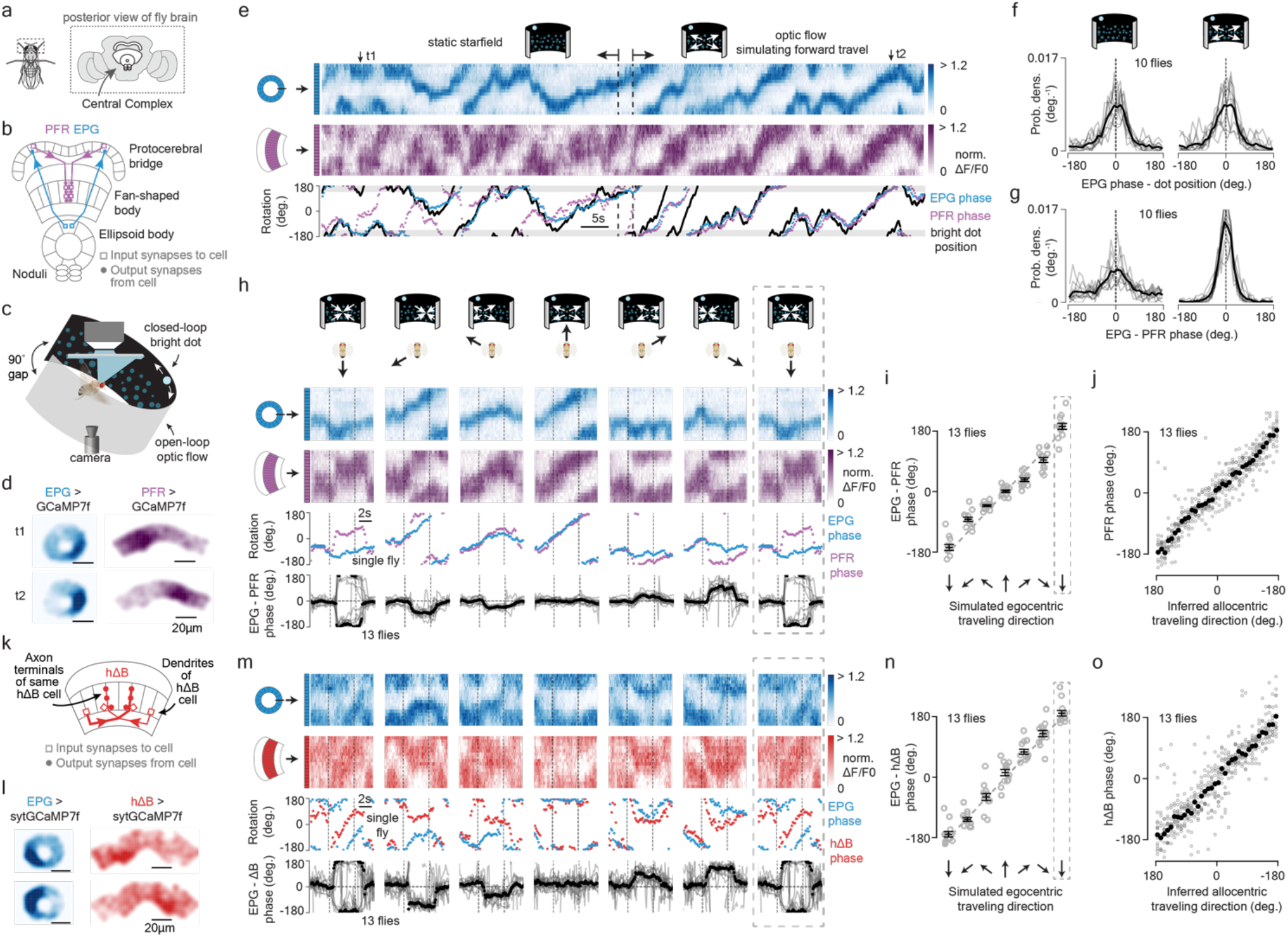
PFR and hΔB neurons signal *Drosophila*’s allocentric traveling direction as differentiated from the allocentric heading signal in EPG neurons. **a**, *Drosophila* brain and central complex. **b**, Close-up of the central complex. Two example EPG cells and two example PFR cells. **c**, Tethered, flying fly setup. **d**, Sample GCaMP7f frames of the EPG bolus in the ellipsoid body and the PFR bolus in the fan-shaped body. **e**, Sample time series of simultaneously imaged EPG and PFR boluses in a tethered, flying fly. Top traces show the [Ca^2+^] signal; bottom traces show [Ca^2+^]-bolus phase estimates and angular position of the closed-loop bright dot. First half: no ventral/lateral optic flow shown. Second half: ventral/lateral optic flow simulating forward travel presented in open loop. Gray regions in bottom trace indicate the 90° rear region of the visual field that contains no LEDs. Time points t1 and t2 in panel d are the same as in this panel. **f**, Probability distributions of the difference between the EPG phase and the bright dot’s angular position, without and with optic flow. Single fly data: light gray. Mean: black. **g**, Same as panel f, but plotting EPG – PFR phase. **h**, Top, EPG (blue) and PFR (purple) GCaMP7f signal in a tethered, flying fly experiencing optic-flow (in the time window bracketed by the vertical dashed lines) with foci of expansion that simulate the following directions of travel: 180° (backward), −120°, −60°, 0° (forward), 60°, 120°, 180° (backward; repeated data). Middle, EPG and PFR phases extracted from the above [Ca^2+^] signals. Bottom, circular-mean phase difference between EPGs and PFRs. Single fly means: light gray. Population mean: black. Dotted rectangle indicates a repeated-data column. **i**, EPG – PFR phase as a function of the egocentric traveling direction simulated by the optic flow. Circular means were calculated in the last 2.5 s of optic flow presentation. Gray: individual fly circular means. Black: population circular mean and s.e.m. Dotted rectangle indicates a repeated-data column. **j**, PFR phase as a function of the inferred allocentric traveling direction, calculated by assuming that the EPG phase indicates allocentric heading direction and adding to this angle, at every sample point, the optic-flow angle. Same data as in panels h-j. Gray: individual fly means. Black: population mean. (Note the flipped x-axis here, which indicates that the PFR bolus position encodes the negative of the fly’s traveling direction; see Methods). **k**, Two example hΔB cells (one thick, one thin) show that the axonal terminals of each hΔB cell are offset from their dendrites by 1/2 the width of the fan-shaped body. **l**, Sample sytGCaMP7f frames of the EPG bolus in the ellipsoid body and the hΔB bolus in the fan-shaped body. **m-o**, Same as panels h-j, but for EPGs and hΔBs.

EPGs represent one of a few dozen sets of *columnar neurons* in the central complex. Each columnar cell class tiles the ellipsoid body, protocerebral bridge, and/or fan-shaped body, and members of each class can be assigned an angular label based on their projection patterns. Many columnar cell classes exhibit [Ca^2+^] boluses that are linked to the EPG bolus and the fly’s heading^15–17^. ***PFRs*** are one such columnar cell class. These cells receive anatomically weak synaptic input from EPGs in the bridge^38^, and they project to the columns of the fan-shaped body where they form extensive, mixed input/output synapses in layers 3, 4 and 5^33,38,39^ (Fig. 1b, purple cells). Imaging [Ca^2+^] in PFRs of both walking (Extended Data Fig. 1) and flying (Fig. 1c, d) flies revealed a single bolus of activity that moves left/right along the fan-shaped body in coordination with the movements of the EPG bolus around the ellipsoid body, consistent with past measurements^17^. Critically, however, we noticed that the PFR and EPG bolus positions sometimes drift apart, especially when walking flies stand still (Extended Data Fig. 1a, b). This raised the possibility that the PFR bolus position might signal the fly’s traveling, rather than heading, angle.

### PFRs and hΔBs signal the fly’s allocentric traveling angle

To test the hypothesis that the PFR bolus position signals the fly’s traveling direction, we focused most of our experiments on tethered, flying flies^40^. In flight, insects rely heavily on panoramic visual motion, or *optic flow*, to assess their direction of travel^41^. We could thus use optic-flow stimuli to systematically simulate the animal’s body translating forward, backward or sideways relative to the heading angle while assessing the effect on the EPG and PFR boluses, measured at the same time via two-photon excitation of GCaMP7f^42^ (Fig. 1c-d). We attached flies to a platform that allowed them to flap their wings in tethered flight and placed the platform in the center of a cylindrical LED arena^40^. A bright dot at the top of the arena rotated in closed-loop with the fly’s steering behavior^40,43^, simulating a static, distant cue that the fly (and the EPG bump) could use to infer heading^14^. A field of dimmer dots in the lower visual field was either stationary or moved coherently to generate open-loop optic-flow that simulated the fly translating along different directions relative to its body axis^44,45^ (Fig. 1c; Methods).

With a stationary dot field (Fig. 1e, left), the EPG phase generally tracked the angular position of the closed-loop dot, consistent with previous studies^14,37^ and with the interpretation that the EPG phase signals the fly’s heading in reference to external cues. The PFR phase, on the other hand, often drifted off the EPG phase, akin to our observations in standing flies. In contrast, when we presented optic flow that simulated the fly’s body moving forward, the EPG and PFR phases became tightly aligned (Fig. 1e, right). Consistent with the sample data in Figure 1e, the EPG phase tracked the closed-loop dot equally well with and without optic flow across ten flies (Fig. 1f), but the PFR phase aligned with the EPG phase much more tightly when optic flow simulated forward translation (Fig. 1g).

When we varied the expansion point of the optic-flow stimulus to simulate the fly translating along six different directions, we observed that the offset between the EPG and PFR phases matched the simulated egocentric (i.e., body-axis-referenced) angle of travel (Fig. 1h, i). If we make the standard assumption that the EPG phase signals the fly’s allocentric (i.e., external-cue-referenced) heading angle^12,14,37^, this implies that the PFR bolus position in the fan-shaped body tracks the fly’s *allocentric traveling direction* (Fig. 1j).

The anatomically dominant input to the PFRs in the fan-shaped body is provided by hDeltaB (***hΔB***) cells^38^, a columnar class in which each constituent cell has a dendritic-input arbor in layer 3 of one fan-shaped body column and a mixed, input/output, axonal arbor in layers 3, 4 and 5 of another column offset by 1/2 the width of the fan-shaped body (Fig. 1k). To image hΔB activity, we created a split-Gal4 driver line to selectively target transgene expression to hΔBs and a UAS-sytGCaMP7f responder line in which GCaMP7f is fused to the c-terminus of synaptotagmin to bias GCaMP to axonal compartments^46^. This bias allowed us to resolve the spatial overlap between the axons and dendrites of different hΔBs and focus on the axonal output of the hΔB population. Imaging sytGCaMP7f fluorescence in hΔBs revealed a bolus of activity at a position along the fan-shaped body that tracks the inferred traveling direction (Fig. 1l-o), much like the PFR bolus. Thus, the traveling direction signal seems to be built first in the hΔBs and then transmitted to the PFRs (as well as, potentially, the many other columnar cell types that hΔBs target^38^).

We also imaged the PFR bolus in tethered, walking flies induced, via optogenetic activation of specific visual-system neurons^47^, to take backward-left or backward-right trajectories on an air-cushioned ball. When flies walked backward, the PFR bolus deviated from the EPG bolus in a manner consistent with the PFR phase signaling the allocentric traveling direction during terrestrial locomotion (Extended Data Fig. 1c-f) as in flight. We also observed deviations of the hΔB phase from the fly’s heading during spontaneous tethered walking behavior in a manner consistent with this signal tracking the fly’s terrestrial traveling direction (data not shown). Thus, the PFR and hΔB boluses appear to track traveling direction in both walking and flying flies.

### PFN_d_s and PFN_v_s are poised to build the traveling signal

To compute their allocentric traveling direction, flies must combine information about their heading angle referenced to external cues (i.e., the EPG heading signal) with information about their egocentric translation direction (i.e., the direction in which they are moving referenced to their body axis). As we will show next, ***PFN***_***d***_ and ***PFN***_***v***_ cells are columnar neurons that receive heading input in the bridge and egocentric translation signals in the noduli^33^ and bridge, while also projecting axons to the fan-shaped body where they synapse onto hΔB neurons^38^ (Fig. 2a, b). PFN_d_s and PFN_v_s thus seem well positioned to convey the EPG heading signal into the fan-shaped body and also to offset that signal when the animal drifts sideward or backward, thereby generating the traveling-direction signal.

**Figure 2.**
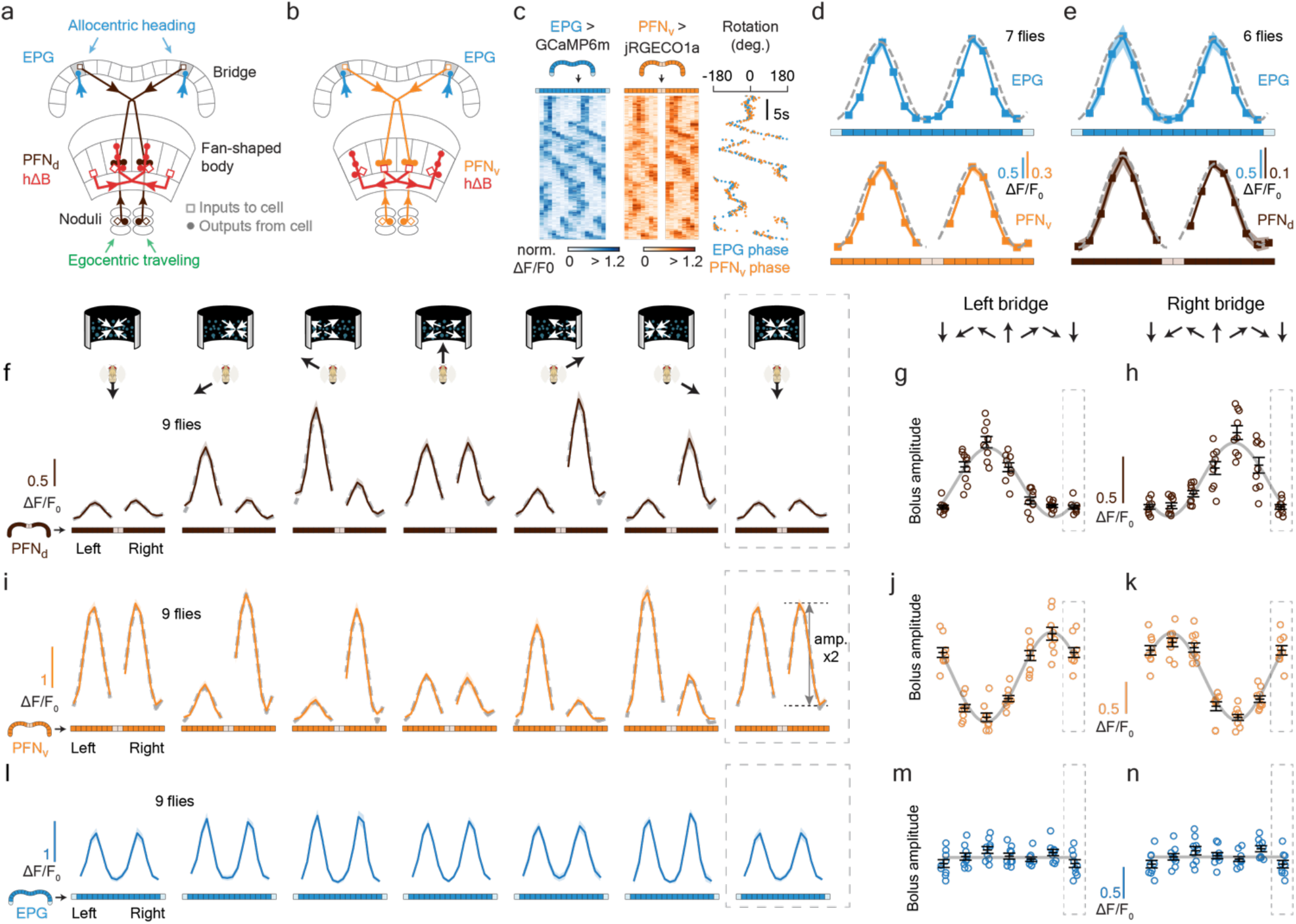
PFN_d_ and PFN_v_ cells show activity patterns consistent with them functioning to build the traveling-direction signal in hΔBs. **a**, EPGs (blue) with dendrites at the top of the ellipsoid body provide direct synaptic input to PFN_d_s (brown) at the two locations highlighted in the bridge, and these PFN_d_s project to the fan shaped body where they synapse on hΔBs. PFN_d_s preferentially target the axon terminals of hΔBs for synapses. PFN_d_s also receive synaptic inputs in the noduli (depicted) and bridge (not depicted), which we found to carry signals related to the fly’s egocentric traveling direction. **b**, EPGs (blue) with dendrites at the top of the ellipsoid body provide direct synaptic input to PFN_v_s (orange) at the two locations shown in the bridge and these PFN_v_s project to the fan shaped body where they synapse on hΔBs. PFN_v_s preferentially target the dendritic region of hΔBs for synapses, which means that PFN_v_s drive an hΔB axonal output signal that is offset from the PFN_v_ input by 1/2 the width of the fan-shaped body. PFN_v_s also receive synaptic inputs in the noduli, which we found to carry signals related to the fly’s egocentric traveling direction. **c**, Sample trace, in tethered flight, of simultaneously imaged GCaMP6m in EPGs and jRGECO1a in PFN_v_s^57^ reveals that the activity boluses of these two cell classes are phase aligned in the bridge. **d**, Phase-nulled EPG (top) and PFN_v_ (bottom) activity patterns in the bridge. Both signals are aligned using the EPG phase. Gray dashed lines: sinusoidal fits. **e**, Same as panel d, but for EPG and PFN_d_ signals in the bridge. **f**, Phase-nulled PFN_d_ activity in the bridge, averaged in the final 2.5 s of the optic flow epoch. Dotted rectangle indicates a repeated-data column, in this panel and throughout. Gray dashed lines: sinusoidal fits. **g**, Single-fly (colored circles) and population means ± s.e.m. (black bars) of the amplitude of PFN_d_ activity profile across the left bridge, averaged in the final 2.5 s of the optic flow epoch. Sinusoidal fit: gray line. **h**, same as panel g, but analyzing right-bridge PFN_d_s. **i-k**, same as panels f-h, but for PFN_v_s. **l-m**, same as panels f-h, but for EPGs. Gray lines in panels m-n: flat-line fits.

Separate arrays of PFN_d_s (Fig. 2a) and PFN_v_s (Fig. 2b) receive monosynaptic input from EPGs in the left and right bridge. Co-imaging EPGs and PFNs in the bridge revealed one activity peak in the left bridge and another in the right bridge for both PFN_d_s and PFN_v_s, with the movement of these peaks closely tracking the movement of the EPG bumps (PFN_v_ and EPG co-imaging: Fig. 2c, d; PFN_d_ and EPG co-imaging: Fig. 2e). Thus, left- and right-bridge PFN_d_s and PFN_v_s together express four copies of the EPG allocentric heading signal. Interestingly, the activity profiles of the four PFN populations across the bridge are sinusoidal (Fig. 2d, e), which is important because, as we will describe below, sinusoidal activity patterns can represent two-dimensional vectors. We hypothesize that the PFN boluses in the bridge are sinusoidally shaped, at least in part, because they receive extensive monosynaptic input from Delta7 **(*Δ7*)** neurons, in the bridge. Δ7s are glutamatergic bridge-interneurons with activity peaks yoked to, but ∼180° offset from, the EPG bumps^48^.

These cells are anatomically and physiologically poised to reshape the EPG bump into a sinusoidally shaped bolus via broad lateral inhibition, not only for PFNs but for a myriad of other columnar cell types within the bridge^38,48^ (Extended Data Fig. 2). Note that the EPGs themselves receive Δ7 feedback, which likely makes their own bridge bumps more sinusoidal than they would otherwise be as well (Fig. 2d-e).

In all our measurements, the PFN boluses were invariably locked to the EPG compass signal, which begs the question: how then does PFN activity also reflect the body’s egocentric translation direction, as would be needed for the PFN activity to build the allocentric traveling signal? We discovered that the amplitudes of the sinusoidal activity patterns of the PFNs across the bridge are strongly modulated by the direction of optic-flow presented to the fly (Fig. 2f-k). Specifically, when we presented optic flow that simulated the fly translating directly forward, the amplitude of the PFN_v_ sinusoids bilaterally dropped, whereas the amplitude of the PFN_d_ sinusoids bilaterally increased. Conversely, when optic-flow simulated backward travel, the amplitude of the PFN_v_ sinusoids bilaterally increased and the amplitude of the PFN_d_ sinusoids bilaterally dropped. With flow that simulated the fly traveling rightward relative to its heading, the amplitude of the right-bridge PFN_d_ sinusoid increased and that of the left-bridge PFN_d_ sinusoid decreased, with the opposite pattern in PFN_v_s. In contrast, the EPG bumps in the bridge show little amplitude modulation with optic-flow (Fig. 2l-n).

Optic-flow modulation of the PFN_v_s most likely arises from the extensive monosynaptic inputs they receive from ***LNO1s*** in the noduli^8,38^. When we imaged activity in the LNO1s, we observed optic-flow tuning that is precisely inverted from that of the PFN_v_s, suggestive of strong, sign-inverting synapses between these cell types (Extended Data Fig. 3). Similarly, PFN_d_s receive extensive monosynaptic inputs from ***LNO2s*** in the noduli as well as from ***SpsPs*** in the bridge^38^. We unfortunately did not have a Gal4 driver line that allowed us to cleanly image LN02s, but when we imaged activity in the SpsPs in the left- and right-bridge, we observed optic-flow tuning that was inverted from the optic-flow tuning observed in PFN_d_s in the bridge, again suggestive of sign-inverting synapses (Extended Data Fig. 3). Although this system is strongly responsive to optic flow in flying flies––where a visual assessment of body translation is critical––we also observed modulations in the activity of LNO1s, SpsPs, PFN_v_s and PFN_d_s in flies walking in complete darkness, consistent with this system employing proprioceptive or efference-copy based estimates of the fly’s egocentric translation direction during terrestrial locomotion in the dark (Extended Data Fig. 3). Because our focus is on the calculation of the traveling direction signal in hΔBs and PFRs in the fan-shaped body, we do not further characterize the nature of the nodulus and bridge inputs to PFNs in this study.

### Combining four sinusoids to build a traveling-direction signal

PFN activity is modulated both by the EPG bump position and the optic-flow drift angle, but how do these modulations drive the hΔB bolus to the appropriate traveling-direction location in the fan-shaped body? Corresponding PFN_v_ and PFN_d_ cells in the left- and right-bridge send projections to the fan-shaped body that are offset from each other by approximately ±1/8 of the extent of the fan-shaped body (Fig. 3a-h). A ±1/8 shift is approximately equivalent to a ±45° angular offset. This ‘push-pull’ anatomy allows the left- and right-bridge PFN_d_s to shift the location of the hΔB activity bolus one way or the other across the fan-shaped body, depending on which has the larger amplitude of activity. PFN_d_s synapse onto both the axonal and dendritic regions of the hΔBs, but, interestingly, the anatomically dominant input is to the axonal region^38^ (Fig. 3a, b). PFN_v_s, on the other hand, also project to the fan-shaped body with the same ±45° angular offset, but they target the hΔB dendrites nearly exclusively (Fig. 3c, d). As described earlier, the axon terminal region of each hΔB is offset from its dendrites by 1/2 the width of the fan-shaped body, equivalent to an angular displacement of ∼180° (Fig. 3a-d). The result of these two sets of shifts is that left- and right-bridge PFN_v_s promote hΔB axonal activity shifted by approximately ±135° relative to their common phase in the bridge. The additional 180° shift in the PFN_v_ pathway compared to the PFN_d_ pathway effectively flips the sign of the PFN_v_ activity sinusoids in the fan-shaped body, thereby compensating for the reversed dependence of the PFN_v_ bolus amplitude on the optic-flow-drift angle relative to the PFN_d_s.

**Figure 3.**
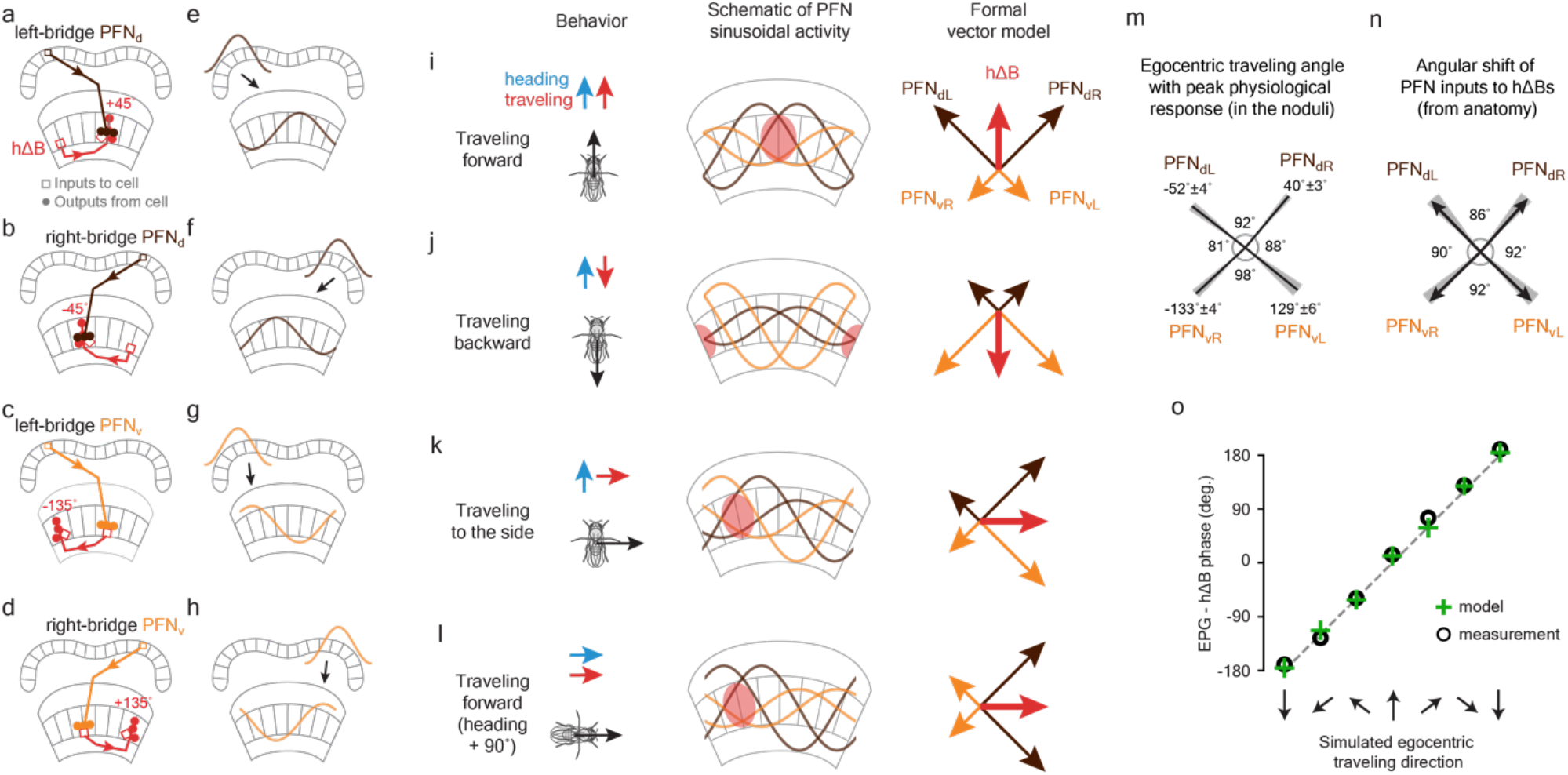
An analytical model of how a vector-sum computation builds a traveling-direction signal in hΔB cells. **a**, Left-bridge PFN_d_s project to the fan-shaped body where they mainly synapse onto the axonal terminals of hΔBs. **b**, Same as panel a, but for the right-bridge PFN_d_s. The corresponding left- and right-bridge PFN_d_ cells are offset from each other by ∼1/4 of the fan-shaped body. **c**, Left-bridge PFN_v_s project to the fan-shaped body where they synapse on the dendritic terminals of hΔBs. **d**, Same as panel c, but for the right-bridge PFN_v_s. Due to the anatomical shift between the axonal and dendritic regions of hΔBs, PFN_v_s and PFN_d_s projecting from the same glomeruli in the bridge induce axonal activity in hΔBs that are offset by ∼1/2 the width of the fan-shaped body. **e-h**, Similar to panels a-d, but showing the sinusoidal activity patterns carried by PFN_d_s and PFN_v_s in the bridge, rather than single cells. In the fan-shaped body, we depict the expected axonal activity pattern in hΔBs, driven by PFNs, based on the anatomy desribed in panels a-d. **i**, When a fly travels forward, both PFN_d_ sinusoids have large amplitude and both PFN_v_ sinusoids have small amplitude, leading the sum of the four vectors, i.e., the hΔB (red) vector, to point forward, or in the heading direction. **j**, When a fly travels backward, both PFN_d_ sinusoids have small amplitude and both PFN_v_ sinusoids have large amplitude, leading the sum, i.e., the hΔB (red) vector, to point backward, or opposite the heading direction. **k**, When a fly travels rightward, the right-bridge PFN_d_ sinusoid and the left-bridge PFN_v_ sinusoid have larger amplitude than their counterparts on the opposite side of the bridge, leading the sum, i.e., the hΔB (red) vector, to point rightward. **l**, Same as panel i––fly walking forward––but after the fly has turned clockwise by 90°, which rotates all the vectors (i.e., the reference frame) by 90° because both the PFN_d_ and PFN_v_ sinusoidal activity patterns have phases that are yoked to the EPG bolus, whose phase rotates with the fly. **m**, The optic-flow-simulated egocentric traveling angle at which the activity of each PFN type is largest depicted with a line at the associated angle. Gray regions: ± s.e.m. Signals are measured in the noduli. Left and right PFN labels refers to the side of the bridge innervated by each cell class, which is opposite to the side of the noduli innervated. **n**, The anatomically calculated angular offset between the four PFN subtypes (black arrows) in how they target the hΔBs (see Extended Data Fig. 5). Gray regions indicate the upper and lower bound of each angle, calculated using different methods (see Extended Data Figs. 4, 5). Note that the sign of angles reported here are flipped relative to those in panels a-d to facilitate the comparison with panel m (see Methods). **o**, There is a good correspondence between the measured deviation of the hΔB phase from the EPG phase (as a function of the optic-flow direction presented) in Figure 1n (circles) and the deviation predicted by a model with no free parameters (plus signs). See Main Text and Methods for details.

These considerations provide a conceptual framework for understanding how an allocentric traveling-direction signal is created in hΔBs. When the fly travels directly forward, both PFN_d_ sinusoids have large amplitudes and both PFN_v_ sinusoids have small amplitudes (Fig. 2f-k). The summed PFN input at the hΔB axon terminals during forward travel will thus peak at the midpoint between the two PFN_d_ boluses, which is the fan-shaped body location that indicates alignment between the fly’s traveling and heading angles (Fig. 3i; the midline of the fan-shaped body for an EPG bump at the top of the ellipsoid body). When the fly travels directly backward, the PFN_d_ and PFN_v_ amplitude modulations switch (Fig. 2f-k), and the summed activity of the four sinusoids yields an activity pattern shifted by 180° (Fig. 3j; 180° offset from the fan-shaped body midline in this example). When the fly translates directly rightward relative to its heading, the left-bridge PFN_v_ sinusoid and right-bridge PFN_d_ sinusoids have larger amplitudes than their counterparts on the opposite side of the bridge (Fig. 2f-k), shifting the summed activity peak by 90° in the fan-shaped body (Fig. 3k; 90° to the left of the midline in this example). This same peak position can be achieved if the fly simply turns 90° to the right, causing all the EPG and PFN signals to rotate 90° in the brain, and then travels straight forward (Fig. 3l). A parallel study reports on a similar set of ideas from data in walking flies^49^.

### A quantitative vector model

We can turn the descriptive framework of the previous paragraph into a quantitative model by noting that adding sinusoidal signals is formally equivalent to adding two-dimensional vectors, which is the basis of an engineering approach known as a *phasor representation*. The fan-shaped body appears to make use of this trick in reverse: to add vectors, it uses the amplitudes and phases of the sinusoidal activity patterns of the PFN_d_ and PFN_v_ populations across the columns of the fan-shaped body to represent the lengths and angles of 2D vectors, and it adds these vectors by summing these neuronal activity profiles. Theoretical models employing phasors have been proposed in studies of the mammalian hippocampus^2,3^ and in insect navigation^6^, including the fan-shaped body^8^, but here we provide the first comprehensive experimental demonstration of their operation.

As already mentioned, the PFN_d_ and PFN_v_ boluses in the bridge are sinusoidally shaped (Fig. 2d-e), consistent with the phasor idea. Moreover, the amplitudes of the four sinusoidal boluses are modulated by optic flow as a cosine function of the drift angle (Fig. 2 g, h, j, k). The maxima of these cosine tuning profiles define the angles of four axes (Figure 3m). According to the map between sinusoids and vectors, the PFN activity profiles represent four vectors with lengths proportional to the projections of the fly’s traveling direction onto these four axes (Methods). For the four vectors to be added properly, the phase differences between the PFN sinusoids in the fan-shaped body––where they are summed by hΔBs––must match the angular difference between the four, physiologically defined axes shown in Figure 3m. The PFN sinusoids in the bridge are (to first order) all aligned with the EPG heading signal, so their phase differences in the fan-shaped body, at the site of hΔB input, are determined by the anatomy of the PFN-to-hΔB projection (Fig. 3a-h). We therefore used the hemibrain connectome to estimate the angular differences of the PFN inputs to hΔBs^38^ (Extended Data Figs. 4, 5; Methods) and, importantly, these differences turned out to be near 90° (Fig. 3n), which is in close agreement with the angular separation across the four axes determined from the PFN response physiology (Fig. 3m). Furthermore, the fact that the phase of all four sinusoids jointly shift with rotations of the EPG compass signal means that the corresponding vectors are referenced to the external world, which implements the ego-to allocentric coordinate transformation. Thus, the four PFN activity sinusoids represent four vectors that, when added, yield a vector that points in the direction of the allocentric traveling direction.

Having established the correspondence between PFN_d_ and PFN_v_ population responses and a set of vectors, we can illustrate, in quantitative detail, how the traveling direction signal is computed (Fig 3i-l, right column). The hΔBs sum the left- and right-bridge PFN_d_ and PFN_v_ activity profiles, which is equivalent to saying that they add the vectors that these neuronal-activity patterns represent, and this vector sum yields the direction of travel. This model can be solved analytically to provide a complete account of the traveling direction computation (Methods). Predictions of the angles encoded by the hΔB bolus made from the model exclusively on the basis of the PFN data points (with no free parameters) are in excellent agreement with the experimental results (Fig. 3o; Methods; see Extended Data Fig. 6 for PFR modeling results).

### Perturbational evidence for the model

To test the vector model, we manipulated PFN_d_ and PFN_v_ activity while measuring the impact on the traveling-direction bolus in PFRs. First, we inhibited EPGs by expressing in them *shibire*^*ts*^, a reagent that abolishes synaptic vesicle recycling at high temperatures^50^. This markedly impairs or eliminates the bolus in EPGs^13^, consistent with the fact that EPGs rely on recurrent connectivity to function normally^15,16,37^. Without EPG input, the PFN_d_ and PFN_v_ sinusoids should be strongly impaired or non-existent. Because the width of the PFR bolus does not notably expand or contract when it is offset from the EPG bolus (analysis not shown), we believe that a recurrent circuit, independent of EPG recurrence, supports a self-sustaining PFR bolus, and thus we expected to observe a PFR bump even in the absence of synaptic input from EPGs and PFNs. However, without effective EPG input to the system, we predicted that the PFR bolus would be unable to tether itself to external cues, let alone track the allocentric traveling angle (Fig. 4a). To stringently test these predictions, we measured the PFR bolus in persistently walking flies, where unlike in flying flies, we rarely observed large deviations of the PFR phase from the angular position of a bright cue or the EPG phase (Extended Data Fig. 1b). With the EPGs silenced, we still observed a clear bolus in PFRs, but the phase of this bolus did not effectively track the angular position of a closed-loop cue (Fig. 4b-d). These data support our prediction that EPG input is necessary for the fan-shaped body traveling signals to be yoked to the external world.

**Figure 4.**
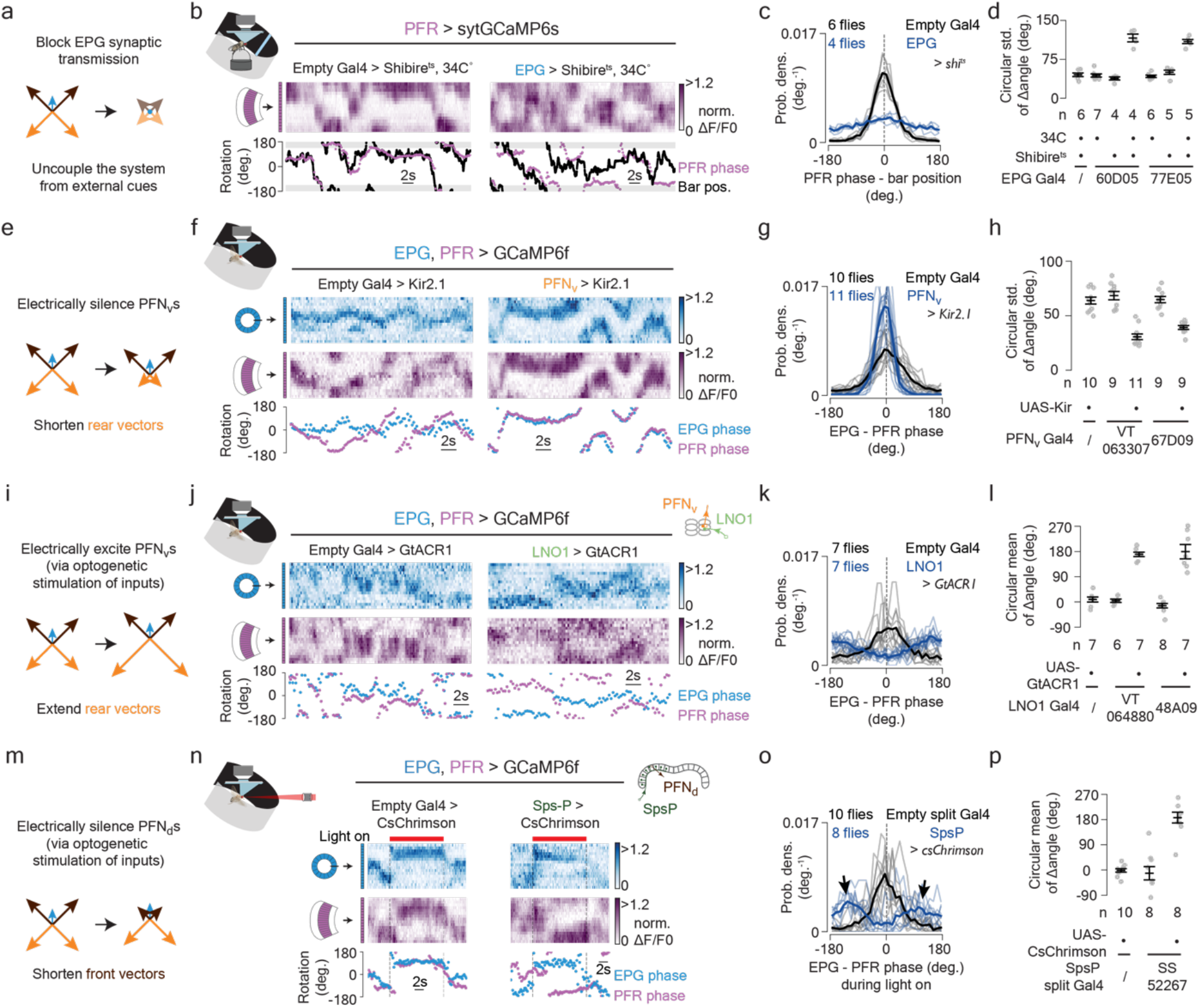
Neural-activity perturbations induce changes in the traveling-direction signal that are consistent with the vector-sum model. **a**, Blocking synaptic transmission in EPGs is akin to diminishing the EPG vector and shortening all four PFN input vectors in the model (since the PFN_d_s and PFN_v_s get EPG and EPG-dependent, Δ7, input). **b**, GCaMP imaging of the PFR bolus in the fan-shaped body of a tethered, walking fly, at 34°C, with EPGs expressing *shibire*^*ts*^ (right) or not expressing *shibire*^*ts*^ (left). **c**, Probability distributions of the difference between the PFR phase and the bright bar’s angular position. Thin lines: single flies. Thick line: population mean. **d**, Circular standard deviation of the PFR – bar position distributions for various genotypes. Gray dots: single fly values. Black dots: population means ± s.e.m. Unpaired t-tests, *P*<2×10^−4^ when comparing the experimental group (4th or 7th columns) with any control group. **e**, Electrically silencing PFN_v_s is akin to shortening the orange, backward pointing vectors. **f**, Simultaneous GCaMP imaging of EPG and PFR activity, in the context of PFN_v_s expressing Kir2.1 (right) or not expressing Kir2.1 (left). **g**, Probability distributions of the difference between the PFR and EPG phase. Thin lines: single flies. Thick line: population mean. **h**, Circular standard deviation of the PFR – EPG phase distributions for various genotypes. Unpaired t-tests, *P*<4×10^−5^ when comparing the two experimental groups (3rd and 5th columns) with any control group. Gray dots: single fly values. Black dots: population means ± s.e.m. **i**, Electrically exciting PFN_v_s (via optogenetic perturbations of their inputs) is akin to lengthening the orange, backward pointing vectors. **j**, Simultaneous GCaMP imaging of PFR and EPG activity, in the context of two-photon-driven, optogenetic silencing of the LNO1 nodulus inputs to PFN_v_s ––which should excite PFN_v_s via a sign-inverting synapse––(right) or when the two-photon activation light was presented in the context of no optogenetic reagent in LNO1s (left). **k**, Probability distributions of the difference between the PFR and EPG phase. Thin lines: single flies. Thick line: population mean. **l**, Circular mean of the PFR – EPG phase distributions for various genotypes. Parametric Watson-Williams multi-sample test, *P*<4⨯10^−4^ when comparing the experimental group (3rd column) with either control group. Gray dots: single fly values. Black dots: population means ± s.e.m. **m**, Electrically exciting PFN_d_s (via optogenetic excitation of their SpsP inputs) is akin to shortening the brown, forward pointing, vectors. **n**, Simultaneous GCaMP imaging of PFR and EPG activity, in the context of two-photon-driven, optogenetic activation of SpsP cells expressing csChrimson (right column). SpsPs synapse heavily on PFN_d_s in the bridge and their activation is expected to silence PFN_d_s via a sign-inverting synapse. A control experiment in which the two-photon activation light was presented in the context of no optogenetic reagent in SpsPs is shown in the left column. **o**, Probability distributions of the difference between the PFR and EPG phase in the context of SpsP activation (blue) and in control flies (gray). Thin lines: single flies. Thick line: population mean. The two black arrows indicate the tendency for the offset to reside at the approximate location of the PFN_v_ sinusoid peaks. **p**, Circular mean of the PFR – EPG phase distributions for various genotypes. Parametric Watson-Williams multi-sample test, P<7×10^−4^ when comparing the experimental group (3rd column) with either control group. Gray dots: single fly values. Black dots: population means ± s.e.m.

Second, we silenced PFN_v_s by expressing in them a K^+^ channel, Kir2.1^51^. This yielded an increase in the phase alignment between the EPG and PFR boluses in tethered, flying flies in the context of no optic flow (Fig. 4e-h). This observation supports our model because silencing PFN_v_s is equivalent to shortening the two, back-pointing, vectors in the model, which physiologically simulates forward motion and should make the PFR vector point in the direction of the EPG angle.

Third, we optogenetically excited PFN_v_s via two-photon activation of the optogenetic Cl^-^ channel, GtACR1^52,53^, in LNO1s^54^, which are the primary monosynaptic, apparently inhibitory (see above and Extended Data Fig. 3), inputs to PFN_v_s in the noduli^38^. Inhibiting LNO1s (and presumably exciting PFN_v_s) drove the PFR bolus to be ∼180° offset from the EPG bolus (Fig. 4i-l). This supports the model because activating PFN_v_s is equivalent to lengthening the two back-pointing vectors in the model, which physiologically simulates backward motion.

Finally, we silenced PFN_d_s by perturbing one of their primary inputs, the SpsPs. There are two SpsP cells per side, each innervating all the ipsilateral PFN_d_s, and the vast majority of SpsP synapses (>80%) in the bridge target PFN_d_s^38^. [Ca^2+^] imaging of SpsPs revealed them to be sensitive to translational optic flow (Extended Data Fig. 3d-f) with tuning inverse to that of the PFN_d_s. We therefore optogenetically activated SpsPs (with csChrimson^55^) to inhibit PFN_d_s. This perturbation drove the PFR bump to be offset by 180°, on average, from the EPG bump (Fig. 4m-p), consistent with the expectation that this perturbation effectively shortens the two front-facing vectors. There appeared to be a tendency for the offset to reside at an angle slightly to the left or right of 180° (Fig. 4o arrows), rather than exactly at 180°, suggesting that the PFR bolus settles to a stable point aligned with one the two PFN_v_ vectors, which remained intact in these experiments.

### Tuning for speed

Thus far we have presented results and discussed the model for translation at a single speed, but we performed some of our experiments on PFNs, hΔBs and PFRs at four different speeds. The model we have presented can account for the PFN responses and the traveling direction angles signaled by the hΔB and PFR boluses at all tested speeds (Extended Data Figs. 7, 8). If the hΔB and PFR boluses were to accurately signal the fly’s traveling vector (angle + speed), rather than just the traveling direction, we would expect the amplitude of their sinusoidal activity profiles to scale with speed. We found that the PFN amplitudes do indeed increase monotonically with speed (Extended Data Fig. 8), but they saturate at high speed and do not vanish at zero speed. This latter feature makes sense if the PFN/hΔB/PFR system aims to ensure the existence of a traveling direction bolus in the fan-shaped body even when flies translate very slowly, but it complicates the representation of a full traveling vector. Overall, PFRs and hΔBs both showed a measurable, but weak, increase in signal strength with faster speeds, but this modulation seems focused to frontal-travel directions (Extended Data Fig. 8 f-g). Speed tuning that is not consistent across different traveling direction angles is incompatible with any simple notion of hΔBs and PFRs carrying a traveling-vector, rather than a traveling-direction, signal. Future work will be needed to relate these varied speed-tuning profiles to their functions in navigation. We suspect that other neurons in the fan-shaped body may show stronger and more consistent speed tuning, which could be useful for building spatial memories via path integration.

## Discussion

Do mammalian brains have neurons whose activity is explicitly tuned to the animal’s allocentric traveling direction as in *Drosophila*? While a defined population of traveling-direction-tuned neurons has yet to be highlighted in mammals, one reason such cells could have been missed is because their activity would loosely resemble that of the head-direction cells outside of a task in which the animal is required to sidestep or walk backwards. Two recent experiments in which rats walked or were transported backwards noted that place cells updated appropriately under these conditions^24,25^. These experiments were conducted in well illuminated environments, but should these same results hold in darkness, where a wholly internal updating of the place cell network is required, angular signals beyond those of head-direction cells, potentially allocentric traveling-direction tuned neurons, would seem to be required to explain the results.

Neurons in area 7a of monkey parietal cortex respond maximally to a specific direction and length of visually driven saccadic eye movements, but the magnitude of this response scales with the starting position of the eye^1^. A classic computational model^1^ demonstrated that such eye-position modulated saccadic responses are expected from the middle layer of a neural network that aims to transform the position of a visual stimulus from the coordinate frame of the retina to that of the head. The *Drosophila* fan-shaped body likewise performs a coordinate transformation from an egocentric sense of body translation, carried by LNO1s and SpsPs (and probably, also, LNO2s), to an allocentric sense, carried by hΔBs and PFRs (Extended Data Fig. 9). In this framework, PFR and hΔB cells develop tuning to the fly’s allocentric traveling direction because neurons at an intermediate processing layer, the PFNs––which show allocentric heading responses that are multiplicatively scaled by the fly’s egocentric translation direction––perform a coordinate transform. Thus, our data provide a biologically validated example of how neurons with *mixed selectivity*, i.e., tuned responses to a primary stimulus axis that are further scaled by a second relevant dimension, function to implement a coordinate transformation, lending credence to the computational ideas from parietal cortex^1,4,5^ and similar proposals in animal navigation^34–36^.

In our model, the optic-flow tuning peaks of left and right PFN_d_s and PFN_v_s (Fig. 3m) are separated from each other by ∼90°, based on measurements made in the noduli (Extended Data Fig. 3i). However, we observed broader separations, close to 120°, when we compared left-right optic-flow tuning in the bridge (Extended Data Fig. 3i). A notable feature of representing and adding vectors as sinusoids is that the system works equally well with non-orthogonal axes. However, if the offset angles in optic-flow tuning are greater than 90°, the anatomical angles describing the shifts in the PFN projections must be correspondingly less than 90° (Methods). An intriguing possibility is that during terrestrial locomotion––where backwards and sideways translation for extended periods is rare––flies may emphasize the egomotion inputs to PFN_d_s found in the bridge and thus employ a non-orthogonal reference frame that may grant them higher spatial resolution in estimating frontal traveling directions. On the other hand, in flight––where backward and sideways translation for extended periods is a distinct possibility––they could rely more on egomotion inputs to PFN_d_s and PFN_v_s in the noduli, employing orthogonal vectors that may allow for better estimating the full 360° of travel accurately. At first, it might seem impossible for the anatomical shifts in the PFN projections to accommodate both of these cases. However, there is an offset in the angular alignment of EPG and Δ7 input to the PFNs in the bridge (Extended Data Fig. 4b, d). Modulating the relative strength of these two inputs will slightly shift the positions of the left- and right-bridge PFN sinusoids, thus changing the angular offsets between sinusoidal inputs to hΔBs in the fan-shaped body and, in principle, allowing a non-orthogonal system to be used in some situations (e.g. walking) and an orthogonal system in others (e.g., flight).

The traveling direction signal we have studied could serve a number of behavioral functions in the fly. For example, it could be compared with a stored, goal direction to generate turning commands to direct the fly along a desired trajectory^13,14^. Also, augmented with an appropriate speed signal, it could provide a basis for path integration, and a model along these lines has been proposed^8^ (see Supplemental Discussion).

Specifically, if one imagines a [Ca^2+^] bolus whose position in the fan-shaped body indicates the traveling direction and whose intensity accurately signals the traveling speed, then a subcellular integration process with a [Ca^2+^]-dependent rate becomes a plausible and attractive mechanism by which instantaneous traveling vectors could be integrated over time to build a memory vector (rather than network-based integration^8^). It has recently been argued that sleep pressure is integrated by a subcellular process in the dorsal fan-shaped body^56^, and this logic might extend to the lower fan-shaped body for integration over shorter––minutes to hours–– timescales related to navigation.

The discovery of an allocentric traveling direction signal is a fundamental step forward for the field of animal navigation. Moreover, because many sensory, motor and cognitive processes can be formalized in the language of linear algebra and vector spaces, defining a neuronal circuit for vector computation could open the door to better understanding a number of previously enigmatic circuits and neuronal-activity patterns across varied nervous systems.

## Acknowledgements

We thank the labs of V. Ruta, B. Dickson and L. Vosshall labs for fly stocks and members of the Maimon laboratory, especially P. Mussells-Pires, for helpful discussions. We thank J. Green and A. Adachi for recombinant stocks, which made this study progress faster than otherwise possible. We thank A. Rizvi for helpful comments on the abstract. We thank Rachel Wilson and Jenny Lu for sharing the hypothesis that the hΔBs may be the locus for the vector integration process rather than PFRs, given anatomical considerations. We thank V. Vijayan and J. Weisman for helping to optimize the csChrimson optogenetics approach during two-photon imaging. We thank A. Kim for sharing code for making optic-flow stimuli, developed off of code initially given to us by P. Weir in the laboratory of M. Dickinson. We thank I. Morantte from the lab of V. Ruta and P. Mussells-Pires for developing the genetic strategy and organizing the plasmids needed to generate the UAS-sytGCaMP7f fly line. Stocks obtained from the Bloomington Drosophila Stock Center (NIH P40OD018537) and the Vienna Drosophila Resource Center were used in this study. Research reported in this publication was supported by a Brain Initiative grant from the National Institute of Neurological Disorders and Stroke (R01NS104934) to G.M. and a Kavli Foundation seed grant to C.L. L.A was supported by NSF NeuroNex Award DBI-1707398 and the Simons Collaboration for the Global Brain. G.M. is a Howard Hughes Medical Institute Investigator.

## Author Contributions

C.L. and G.M. conceived of the project. C.L. performed the experiments and analyzed the data. C.L., G.M., and L.A. jointly interpreted the data and decided on new experiments. L.A. developed and implemented the formal models. C.L. wrote the initial draft of the paper, which was then edited by G.M. and L.A.

## Author Information

The authors declare no competing financial interests.

## Supplemental Discussion

### Angular integration and vector addition in the fly central complex

PFNs project from the bridge to the fan-shaped body in an anatomically offset manner compared to how PFRs project from the bridge to the fan-shaped body; this pattern is very reminiscent of how PENs project from the bridge to the ellipsoid body in an offset manner to EPGs^33^ (Extended Data Fig.10). The PEN/EPG circuit implements angular integration^15,16^, which means that a persistent asymmetry in the activity of left-vs. right-bridge PENs––induced when the fly is turning––yields a *persistent rotation* of the EPG bolus. We argue here that the PFN circuit implements vector addition, which means that a persistent asymmetry in the activity of left-vs. right-bridge PFNs––induced when the fly is translating left or right––yields a *persistent offset* in the PFR bolus relative to the EPG bolus rather than a persistent rotation. The EPG phase continuously rotates when there is a constant asymmetric input because the EPG system contains functionally relevant projections both from the bridge to the ellipsoid body and from the ellipsoid body back to the bridge. This closed-loop anatomy allows the EPG bump to rotate indefinitely with proper input drive. In the PFN/hΔB/PFR system, there are only unidirectional projections from the bridge to the fan-shaped body, which means that a constant asymmetric input only nudges the system, in open loop, to an offset position, rather than inducing a continuous precession. Thus, our work highlights how very similar anatomical relationships between cells, in the PEN/EPG versus PFN/PFR systems, can yield quite different circuit-level computations.

### Relating our results to a model of path integration in the bee fan-shaped body

A previously proposed computational model of path integration, based on anatomical and physiological experiments in bees^8^, bears some relation to our findings in *Drosophila*. For example, the basic observation that noduli inputs to PFNs have optic-flow tuning curves with peaks skewed to the left and right of the fly’s midline was noted by these authors, and it was presciently proposed that these inputs might allow bees to estimate their direction of travel relative to their body axis. However, the authors did not propose the existence of an explicit, traveling-direction bolus in the fan-shaped body; rather, the traveling velocity signal they extracted was carried by a difference between left- and right-projecting cell populations. These authors further hypothesized that PFNs, or “working-memory CPU4 cells”, integrate a speed-modulated traveling-direction signal over time in the PFN network itself (perhaps via recurrent PFN-to-PFN connections) to generate a persistent memory of the direction and distance from a start location. Our work does not reveal the PFN_v_s and PFN_d_s to be integrating their bridge and noduli inputs over time. Rather, the PFN_d_s and PFN_v_s appear to adjust the position of the hΔB and PFR [Ca^2+^] bumps in the fan-shaped body in real-time. That said, beyond PFN_d_s and PFN_v_s, which innervate the third layer of the fan-shaped body, there exist three additional classes of PFNs in layers one and two^33^, and the activity of these PFNs may resemble the bee model more closely. Furthermore, it is possible, although we think it unlikely, that PFN_v_s and PFN_d_s switch to performing a more memory-integration function in the context of flies performing a path-integration task, an idea that could be tested in future work.

### Estimating the traveling direction with optic flow is robust to changes in the head angle

When a fly flies forward with its head aligned to its body axis, it experiences optic flow with a focus of expansion that is centered along the visual field’s midline. If the fly were to rotate its head to the right by, say, 20°, while still flying forward, this would shift the optic-flow’s focus of expansion, and the simulated egocentric traveling direction, by 20° to the left, which would seem to introduce an error in the traveling-direction calculation. Importantly, however, the same head movement would also rotate the perceived angle of distant visual cues (like the sun) by the exact same amount, and the EPG system would register this as a 20° rotation as well. (The same logic would hold for non-visual cues sensed in the head’s reference frame and relevant for the EPG heading signal, like the antennal sensing of wind direction.) The 20° leftward rotation of the egocentric traveling angle would cancel the 20° rightward rotation in the EPG’s allocentric heading estimate to yield, notably, the same allocentric traveling angle in hΔBs or PFRs (Extended Data Fig. 11). In this way, our circuit naturally compensates for varying yaw head angles during flight. This logic assumes and perhaps even intimates that the EPG’s bump position signals the fly’s head angle rather than the body angle. The same logic would not hold if the fly were relying on proprioceptive (or efference-copy) signals from the leg motor-system (rather than optic flow signals from the eyes) to estimate the egocentric traveling direction, which is likely to be the case in walking. In this situation a coordinate transformation would be needed to place egocentric, traveling-direction signals (from the body) in the same reference frame as allocentric, head-direction signals (from the eyes), before impacting the hΔB/PFR boluses in the fan-shaped body.

## Methods

### Fly husbandry

Flies were raised at 25°C with a 12-h light and 12-h dark cycle. In all experiments, we studied female *Drosophila melanogaster* that were 2-6 days old. Flies were randomly selected for all of experiments. We excluded flies that appeared unhealthy at the time of tethering as well as flies that did not fly longer than 20 s in flight experiments. This meant excluding fewer than 5% of flies for most genotypes. However, in the perturbational experiments shown in Figs. 4e-p many flies flew poorly––perhaps because these genotypes all expressed 5-6 transgenes which can affect overall health and flight vigor––and we had to exclude ∼70% of flies due to poor tethered flight behavior (i.e., would not maintain continuous flight for > 5 s bouts). The 30% of flies tested in these genotypes flew in bouts that ranged 20 s to many minutes, allowing us to make the necessary EPG/PFR signal comparisons (discussed below). Flies in optogenetic experiments were shielded from green and red lights during rearing by placing the fly vials in a box with blue gel filters (Tokyo Blue, Rosco) on the walls. After eclosion, two days or more prior to experiments, we transferred these flies to vials with food that contained 400µM all-*trans*-retinal.

### Cell type acronym naming conventions

| Names in this paper<br>(in hemibrain v1.1 if<br>different) | Names<br>used in<br>reference <sup>54</sup> | Description of acronym | References in which cell type<br>is studied or defined |
| --- | --- | --- | --- |
| EPG | E-PG | Ellipsoid body-Protocerebral<br>bridge-Gall | Lin C, et al<sup>39</sup>; Wolff T, et<br>al<sup>33,54</sup>; Seelig J, et al<sup>12</sup>; Green J,<br>et al<sup>15</sup>; Turner-Evans D, et al<sup>16</sup> |
| PFR (PFR_a) | P-F-R | Protocerebral bridge-Fan shaped<br>body- Fan shaped body-Round<br>body | Lin C, et al<sup>39</sup>; Wolff T, et<br>al<sup>33,54</sup>; Shiozaki H, et al<sup>17</sup> |
| PFN<sub>v</sub> (PFN<sub>v</sub>) | P-FN<sub>v</sub> | Protocerebral bridge-Fan shaped<br>body- Nodus (ventral) | Lin C, et al<sup>39</sup>; Wolff T, et<br>al<sup>33,54</sup> |
| PFN<sub>d</sub> (PFN<sub>d</sub>) | P-FN<sub>D</sub> | Protocerebral bridge-Fan shaped<br>body- Nodus (dorsal) | Lin C, et al<sup>39</sup>; Wolff T, et<br>al<sup>33,54</sup> |
| hΔB (hDeltaB) | -- | Columnar cell class with lateral<br>projections in layer 3 of the fan-shaped body | -- |
| LNO1 | L-N | Lateral accessory lobe-Nodus | Wolff T, et al<sup>54</sup> |
| LNO2 | -- | Lateral accessory lobe-Nodus | -- |
| SpsP | Sps-P | Superior posterior slope-Protocerebral bridge | Wolff T, et al<sup>33,54</sup> |
| Δ7 (Delta7) | Delta7 | Skiping 7 glomeruli in the<br>Protocerebral bridge between two<br>output areas | Lin C, et al<sup>39</sup>; Wolff T, et<br>al<sup>33,54</sup>; Turner-Evans D, et al<sup>48</sup> |
| PEN1 (PEN_a) | P-EN1 | Protocerebral bridge-Ellipsoid<br>body - Nodus | Lin C, et al<sup>39</sup>; Wolff T, et<br>al<sup>33,54</sup>; Green J, et al<sup>15</sup>; Turner-Evans D, et al<sup>16</sup> |
| PEN2 (PEN_b) | P-EN2 | Protocerebral bridge-Ellipsoid<br>body - Nodus | Green J, et al<sup>15</sup> |

#### Fly genotypes

- For simultaneous imaging of EPGs and PFRs:

In Fig. 1e-j and Extended Data Figs. 6, 7, we used + (Canton S, Heisenberg Laboratory); UAS-GCaMP7f (Bloomington Drosophila Stock Center, BDSC #80906)/+; 60D05-Gal4 (BDSC #39247)/37G12-Gal4 (BDSC #49967) flies.

In Fig. 4f-h, we used +; 60D05-LexA, LexAop-GCaMP6f (BDSC#44277)/37G12-LexA (BDSC #52765); UAS-Kir2.1 (Leslie Vosshall Laboratory)/VT063307-Gal4 (Vienna Drosophila Resource Center, CDRC) flies and +; 60D05-LexA, LexAop-GCaMP6f/37G12-LexA; UAS-Kir2.1/67D09-Gal4 (BDSC #49618) flies for the experimental groups. For the control groups, we used +; 60D05-LexA, LexAop-GCaMP6f/37G12-LexA; UAS-Kir2.1/empty-Gal4 (BDSC #68384) flies, +; 60D05-LexA, LexAop-GCaMP6f/37G12-LexA; VT063307-Gal4/empty-Gal4 flies, and +; 60D05-LexA, LexAop-GCaMP6f/37G12-LexA; 67D09-Gal4/empty-Gal4 flies.

In Fig. 4j-l, we used +; 60D05-LexA, LexAop-GCaMP6f /37G12-LexA; UAS-GtACR1-EYFP (Adam Claridge-Chang Laboratory)/VT064880-Gal4 (CDRC) flies and +; 60D05-LexA, LexAop-GCaMP6f /37G12-LexA; UAS-GtACR1-EYFP/48A09-Gal4 (BDSC #50342) flies for the experimental groups. For the control groups, we used +; 60D05-LexA, LexAop-GCaMP6f/37G12-LexA; UAS-GtACR1-EYFP/empty-Gal4 flies, +; 60D05-LexA, LexAop-GCaMP6f/37G12-LexA; VT064880-Gal4/empty-Gal4 flies, and +; 60D05-LexA, LexAop-GCaMP6f/37G12-LexA; 48A09-Gal4/empty-Gal4 flies.

In Fig. 4n-p, we used UAS-CsChrimson-mVenus (BDSC #55134)/+; 60D05-LexA, LexAop-GCaMP6f/ VT019012-AD; VT005534-LexA (Barry Dickson Laboratory)/72C10-DBD (Janelia FlyLight Split-Gal4 Driver Collection, FlyLight SS52267) flies for the experimental groups. For the control groups, we used UAS-CsChrimson-mVenus; 60D05-LexA, LexAop-GCaMP6f/empty-LexA (BDSC #77691); VT005534-LexA/empty-Gal4 flies, and + (Canton-S); 60D05-LexA, LexAop-GCaMP6f/ VT019012-AD; VT005534-LexA (Barry Dickson Laboratory)/72C10-DBD flies.

In Extended Data Fig. 1b, we used +; UAS-GCaMP6m (BDSC #42748)/+; 60D05-Gal4 /37G12-Gal4 flies.

In Extended Data Fig. 1d-f, we used UAS-CsChrimson-mVenus/+; 60D05-LexA, LexAop-GCaMP6f/ 26A03-AD; VT005534-LexA/54A05-DBD (FlyLight OL0046B) flies.

In Extended Data Fig. 2h (bottom row), we used +; 60D05-LexA/+; LexAop-GCaMP6m, UAS–jRGECO1a (BDSC #63794)/37G12-Gal4 flies.

- For simultaneous imaging of EPGs and hΔBs:

In Figs. 1l-o and 3o, we used +; UAS-sytGCaMP7f /72B05-AD (BDSC # 70939); 60D05-Gal4/ VT055827-DBD (BDSC # 71851) flies. We created the sytGCaMP7f construct by linking the GCaMP7f and *Drosophila* synaptotagmin 1 coding sequences using a 33GS linker. We then used this construct to generate transgenic flies by PhiC31-based integration into attp40 site, performed by BestGene.

- For simultaneous imaging of EPGs and PFN_d_s:

In Fig. 2e and Extended Data Fig. 2h (top row), we used +; 60D05-LexA/+; LexAop-GCaMP6m, UAS– jRGECO1a /47E04-Gal4(BDSC #50311) flies.

- For simultaneous imaging of EPGs and PFN_v_s:

In Fig. 2c, d and Extended Data Fig. 2e (left column), h (middle row), we used +; 60D05-LexA/+; LexAop-GCaMP6m, UAS–jRGECO1a/VT063307-Gal4 flies.

- For imaging single cell types:

In Fig. 2f-h and Extended Data Figs. 3e-i, n, o, 4g, i, 8a-c, h, i, n, o, we used +; 15E01-AD (BDSC #70558)/+; UAS-GCaMP7f (BDSC #79031)/47E04-DBD (BDSC #70366) flies for PFN_d_s.

In Fig. 2i-k and Extended Data Figs. 3b, c, i, j, k, 4h, i, 8d, e, j, k, p, q, we used +; +; UAS-GCaMP7f/VT066307-Gal4 flies for PFN_v_s.

In Fig. 2l-n and Extended Data Fig. 4e, f, we used +; +; UAS-GCaMP7f/60D05-Gal4 flies for EPGs.

In Fig. 4b-d, we used pJFRC99-20XUAS-IVS-Syn21-Shibire-ts1-p10 inserted at VK00005 (referred to here as UAS-*shibire*^*ts*^) to drive *shibire*^*ts*^ (Rubin Laboratory). We used +; 37G12-LexA/LexAop-sytGCaMP6s (Vanessa Ruta Laboratory); 60D05-Gal4/UAS-*shibire*^*ts*^ flies for the experimental group. For the control groups, we used +; 37G12-LexA/LexAop-sytGCaMP6s; 60D05-Gal4/empty-Gal4 flies and +; 37G12-LexA/LexAop-sytGCaMP6s; UAS-*shibire*^*ts*^/empty-Gal4 flies.

In Extended Data Fig. 3b, c, i, l, m, we used +; VT020742-AD /+; UAS-GCaMP7f/VT017270-DBD (FlyLight SS47398) flies for LNO1s.

In Extended Data Fig. 3e, f, i, p, q, we used +; VT019012-AD/+; UAS-GCaMP7f/72C10-DBD (FlyLight SS52267) flies for SpsPs.

In Extended Data Fig. 8f, l, r, we used +; UAS-sytGCaMP7f /72B05-AD; +/ VT055827-DBD flies for hΔBs. In Extended Data Fig. 8g, m, s, we used +; +; UAS-GCaMP7f/37G12-Gal4 flies for PFRs.

### Distinguishing PFR subtypes in Gal4 lines

The hemibrain connectome^38^ defines two subtypes of PFR cells^33^: PFR_a cells and PFR_b cells, which differ in the details of their projections and connectivity in the fan-shaped body. Both PFR_as and PFR_bs are columnar cells that project from the protocerebral bridge to the fan-shaped body. Based on the connectome^38^, PFR_as and PFR_bs that innervate 4 of the bridge glomeruli project to the fan-shaped body in the same way and PFR_as and PFR_bs that innervate 12 other bridge glomeruli project to the fan-shaped body in a slightly different way. We used this fact to interrogate the MultiColor Flip-Out (MCFO) single-cell anatomical data set^58^ from the FlyLight Generation 1 MCFO Collection to quantify the ratio of each PFR subtype in the two Gal4 driver lines that we used for targeting transgenes to PFRs. For driver line 37G12, we found 10 out of 13 cells in the MCFO data had an innervation pattern that is consistent with PFR_a but not PFR_b and the innervation patterns of the other 3 cells were indistinguishable between the two subtypes. For driver line VT005534, we found 2 out of 3 cells in the MCFO data were consistent with them being PFR_as and not PFR_bs and the third cell had an anatomy that did not allow us to distinguish between subtypes. We observed no cell whose projection pattern matched PFR_b but not PFR_a. These results argue that the majority of the PFR cells targeted by the two Gal4 driver lines we used are PFR_a cells.

### Fly preparation and behavioral setup

As described previously, we glued flies to a custom stage for imaging during flight^40^ and to a slightly different custom stage––which allows for more emission light to be collected by the objective––for imaging during walking^15^. Dissection and imaging protocols followed previous studies^15^. For tethered flight experiments, each fly was illuminated with 850 nm LEDs with two fiber optics from behind^40^. A Prosilica GE680 camera attached to a fixed-focus Infinistix lens (94-mm working distance, 1.0× magnification, Infinity) imaged the fly’s wing-stroke envelope at 80-100 Hz. The lens also held an OD4 875-nm shortpass filter (Edmund Optics) to block the two-photon excitation laser (925 nm). This camera was connected to a computer that tracked the fly’s left and right wing beat amplitude with custom software developed by Andrew Straw (https://github.com/motmot/strokelitude)^40^. Two analog voltages were output in real time by this software and the difference between the left and right wingbeat amplitude was used to control the angular position of the bright dot on the LED arena in closed-loop experiments (described below). For tethered walking experiments we followed protocols described previously^15^.

### LED arena and visual stimuli

We used a cylindrical LED arena display system^59^ with blue (465 nm) LEDs (BM-10B88MD, Betlux Electronics). The arena was 81° high and wrapped around 270° of the azimuth, with each pixel subtending ∼1.875°. To minimize blue light from the LEDs inducing noise in the microscope’s photomultiplier tubes, we reduced the LED intensities, over most of the arena, by covering the LEDs with five sheets of blue gel (Tokyo Blue, Rosco). Over the top 16 pixels top of the arena, we only placed two gel sheets, so that the closed-loop dot at the top of the arena was brighter than the optic flow at the bottom, which may have helped to promote that the fly interpret the bright blue dot as a celestial cue (like the sun) and the optic-flow at the bottom as ground or side motion. During flight experiments, we held the arena in a ∼66° pitched-back position, so that the LED vertical and horizontal axes matched the major ommatidial axes of the eye^45^. During walking experiments, we typically presented a tall vertical bar––rather than a small dot––in closed loop and we tilted the arena by only ∼30° because the ball physically occludes the ventral visual field and a shallower arena tilt made it more likely that the fly could see the closed-loop stimulus over all 270° of the azimuthal positions that it could take.

We adapted past approaches for generating optic flow (starfield) stimuli^44,45^. In brief, we populated a virtual 3D world with forty-five, randomly positioned spheres (2.3-cm diameter) per cubic meter. The spheres were bright on a dark background. We only rendered spheres that were within two meters of the fly because spheres further away contributed only minimally to the observed motion and, if rendered, would have overpopulated the visual field with bright pixels. We then calculated the angular projection of each sphere onto the fly’s head and used this projection to determine the pattern to display on the LED arena on each frame. To prevent the size of each sphere from being infinitely large as it approached the fly, we limited each sphere’s diameter on the arena to be no larger than 7.5°. The starfield extended from 4 pixels (∼8°) above to 20 pixels (∼40°) below the arena’s midline. In all experiments that employed open-loop optic flow (Figs. 1, 2f-n, 3o and Extended Data Figs. 3b-i, 4, 6e, 7, 8), the optic-flow position was updated at a frame rate of 25 Hz. (Note that the LED *refresh* rate was at least 372 Hz^59^.) To simulate the optic flow that a fly would experience when it is translating through 3D space, we moved a virtual fly in the desired direction(s) through the virtual world and displayed the resultant optic flow pattern on the arena. We used a translation speed of 35 cm/s in all experiments, except those in Extended Data Figures 7 and 8 where we tested multiple speeds as indicated (ranging from 8.75 to 70 cm/s). Although we report on the translation speed of the virtual fly in metric units, the optic flow experienced by insects translating at 35 cm/s will vary dramatically depending on the clutter of the local environment. We believe that the optic flow stimuli we presented in our study are likely to be in an ethologically relevant range because (1) our virtual fly translated at speeds that bracket observed flight speeds in natural environments^60– 62^ and (2) optic-flow sensitive cells reported on here (and others not discussed in this study) responded with progressively increasing activity to the presented stimuli/speeds rather than immediately saturating or showing no detectable responses (e.g., Extended Data Fig. 8). That said, our stimuli simulated a dense visual environment and it will be important to test our results in the context of reduced visual clutter in future work.

In flight experiments with a closed-loop dot (Figs. 1, 2c-e, 3o and Extended Data Figs. 2, 3b-f, i, 6e, 7), the dot subtended 3.75° by 3.75° and was located ∼34° above the arena’s midline. We used the difference of the left and right wingbeat amplitudes (L–R WBA) to control the azimuthal velocity of the bright dot on the LED arena. That is, when the right wingbeat amplitude is smaller than the left wingbeat amplitude (indicating that the fly is attempting to turn to the right), the dot rotated to the left, and vice versa. The negative-feedback closed-loop gain was set to 7.3°/s per degree change in L–R WBA. In closed-loop walking experiments with a visual stimulus (Fig. 4b-d and Extended Data Fig. 1), the bright bar was 11.25° wide and spanned the entire height of the arena. We directly linked the azimuthal position of the bright bar on the LED arena to the azimuthal position of the ball under the fly using Fictrac^15,63^, as described previously. This closed-loop set up mimics the visual experience of a fly with a bright cue at visual infinity, like the sun. We did not provide translational stimuli in closed loop in this paper.

### Calcium imaging

We used a two-photon microscope with a movable objective (Ultima IV, Bruker). The two-photon laser (Chameleon Ultra II Ti:Sapphire, Coherent) was tuned to 1000-1010 nm for simultaneous imaging of GCaMP6m and jRGECO1a (Fig. 2c-e and Extended Data Fig. 2), and was otherwise tuned to 925 nm in all of the other imaging experiments. We used a 40x/0.8 NA objective (Olympus) or 16x/0.8 NA objective (Nikon) for all imaging experiments. The laser intensity at the back aperture was 30-40mW for walking experiments and 40-80mW for flight experiments. Because of light loss through the objective and the fact that the platform to which the fly was attached blocks ∼half the light from reaching the fly, we estimate an illumination intensity, at the fly, of ∼16-32mW for flight experiments. In walking experiments, the platform to which we attach the fly blocks less light and we expect an illumination intensity, at the fly, of ∼24-32mW. A 575 nm dichroic split the emission light. A 490-560nm bandpass filter (Chroma) was used for the green channel PMT and a 590-650nm bandpass filter (Chroma) was used for the red channel PMT. We recorded all imaging data using three to five z-slices, with a Piezo objective mover (Bruker Ext. Range Piezo), at a volumetric rate of 4-10 Hz. We perfused the brain with extracellular saline composed of (in mM) 103 NaCl, 3 KCl, 5 N-Tris(hydroxymethyl) methyl-2-aminoethanesulfonic acid (TES), 10 trehalose, 10 glucose, 2 sucrose, 26 NaHCO3, 1 NaH2PO4, 1.5 CaCl2, 4 MgCl2, and bubbled with 95% O2/5% CO2. The saline had a pH of ∼7.3 and an osmolarity of ∼280 mOsm. We controlled the temperature of the bath by flowing the saline through a Peltier device and measured the temperature of the bath with a thermistor (Warner Instruments, CL-100).

### Optogenetic stimulation

In the optogenetic experiments in Fig. 4j-l, we used the two-photon laser tuned to 925 nm to excite GtACR1, with the same scanning light being used to excite GCaMP. To excite CsChrimson in the optogenetic experiments in Fig. 4n-p and Extended Data Fig. 1), we focused 617nm laser (M617F2, Thorlabs) on to the front, middle of the fly’s head with a custom lens set (M15L01 and MAP10100100-A, Thorlabs). We placed two bandpass filters (et620/60m, Chroma) in the two-photon microscope’s emission path to minimize any of the optogenetic light being measured by the photomultiplier tubes. In flight experiments (Fig. 4n-p), we used pulse-width modulation at 490Hz (Arduino Mega board) with a duty cycle of 0.8 to change the 617 nm laser’s intensity. We measured the laser’s intensity at the fly’s head to be 20.8 µW. In the experiments where we triggered backward walking via activation of csChrimson (Extended Data Fig. 1) in lobula columnar neurons, the duty cycle of the red light was 0.7 and the effective light intensity was 18.2 µW.

In Fig. 4j-l, for two main reasons, rather than directly exciting PFN_v_s, we optogenetically inhibited the LNO1 inputs to PFN_v_s. First, [Ca^2+^] imaging revealed opposite responses to our optic flow stimuli in the two cell types (Extended Data Fig. 3a-c), as mentioned earlier, arguing for a sign-inverting synapse between them. Second, we tried optogenetically activating the PFN_v_s directly (data not shown), which yielded more variable movements of the PFR bolus. We believe that stimulating LNO1s yielded more consistent effects on the PFR bolus because there are only two LNO1s per side and they synapse uniformly on all PFN_v_s within a tiny neuropil (the second layer of the nodulus) on their side^38^. Stimulating GtACR1 in a small volume likely made homogeneous activation of the PFN_v_ population more feasible.

### Data analysis

#### Data acquisition and alignment

All data were digitized by a Digidata 1440 (Molecular Devices) at 10kHz, except for the two-photon images which were acquired using PrairieView (Bruker) at varying frequencies and saved as tiff files for later analysis. We used the frame triggers associated our imaging frames (from Prairie View), recorded on the Digidata 1440, to carefully align behavioral measurements with [Ca^2+^] imaging measurements.

#### Experimental Structure

For Fig. 1d-g, each fly performed tethered flight while in control of a bright dot in closed loop. Each recording was split into three segments, where we first presented a static starfield for 90 s, followed by progressive optic flow for 90 s, and ending with another static starfield for 90 s.

For Fig. 1h-j and Extended Data Fig. 6e, we presented each fly with a closed-loop dot throughout. We presented four blocks of eight optic-flow stimuli (six translational plus two rotational) per block, shown in a pseudorandom order. Each 4 s optic flow stimulus was preceded and followed by 4 s of optic flow that mimics forward travel, which ensured that the EPG and PFR bumps were aligned––for a stable “baseline”––before and after each tested optic flow stimulus. We presented 4 s of a static starfield between each repetition of the above three patterns. We employed the same protocol for Figs. 1l-o and 3o, but we just presented the six translational optic flow stimuli, without the two, rotational optic flow stimuli.

For Fig. 2c-e and Extended Data Fig. 2e, h, each fly was presented with a closed-loop dot and static starfield throughout the recording. Recording durations ranged from 1-4 min.

For Figs. 2f-n and Extended Data Figs. 3e-h (PFN_d_ rows), i (PFN_d_s in the noduli, PFN_d_s and PFN_v_s in the bridge), 4, 8, we did not have a closed-loop dot. We presented four blocks of 24 stimuli (6 translational directions at 4 different speeds) per block, shown in a pseudorandom order. Each stimulus was preceded by 1.2-s of a static starfield, followed by 4 s of optic flow at different directions, and ending with a 1.2-s static starfield.

For Fig. 4b-d, each fly was presented with a tall bright bar in closed-loop throughout. EPG > Shibire^ts^ flies experienced both 25°C and 34°C trials in these experiments and we waited ∼5 min after the bath temperature reached 34°C before imaging in the EPG silenced condition so as to increase the likelihood of thorough vesicle depletion. Recording durations ranged from 6-8 min.

For Fig. 4f-h, j-l, flies performed tethered flight in the context of a dark (unlit) visual display. We recorded data for ∼1-4 min and if the fly was flying robustly, we collected a second data set from the same fly.

For Fig. 4n-p, flies performed tethered flight in the context of a dark (unlit) visual display. We recorded data for 2-6 min. and if the fly was flying robustly, we collected a second data set from the same fly. We presented 12 s red light pulses to activate csChrimson every ∼20 s.

For Extended Data Fig. 1b, each fly was presented with a tall bright bar in closed-loop throughout. We recorded data for 10 min., with up to three 10-min. data sets collected per fly.

For Extended Data Fig. 1c-f, each fly was presented with a tall bright bar in closed-loop throughout. We recorded data for 7-10 min., with up to 3 data sets collected per fly. We presented 4 s red light pulses to activate csChrimson every 1-3 min.

For Extended Data Fig. 3b, c, e-f (SpsP rows) and i (PFN_v_s in the noduli, SpsPs and LNO1s), we presented each fly with a closed-loop dot throughout. We presented four blocks of eight optic-flow stimuli (six translational plus two rotational) per block, shown in a pseudorandom order. Each optic flow stimulus was preceded by 4-s of a static starfield, followed by 4-s of optic flow at different directions, and ending with a 4-s of static starfield.

For Extended Data Fig. 3j-q, each fly was walking in the dark. We recorded data for 5 min., with up to three 10-min. data sets collected per fly.

For Extended Data Fig. 7, we presented each fly with a closed-loop dot throughout. We presented three blocks of 18 stimuli per block (6 translational directions at 3 different speeds), shown in a pseudorandom order. Each stimulus was preceded by 1.2-s of a static starfield and 4 s of translational optic flow simulating forward travel, followed by 4 s of optic flow at different directions, and ending with a 4 s of optic flow simulating forward travel.

### Image registration

Before quantifying fluorescence intensities, imaging frames were registered in Python by translating each frame in the x and y plane to best match the time-averaged frame for each z-plane. Multiple recordings from the same fly were registered to the same time-averaged template if the positional shift between recordings was small.

### Defining regions of interest

To analyze calcium imaging data, we defined regions of interest (ROIs) in Fiji and Python for each glomerulus (protocerebral bridge), wedge (ellipsoid body) or column (fan shaped body). For the bridge data, we defined ROIs by manually delineating each glomerulus from the registered time-averaged image of each z-plane (Fig. 2 and Extended Data Figs. 2, 3e-h PFN_d_ row, i PFN_d_s and PFN_v_s in the bridge, j-k, n-o, 4, 8a-e, h-k, n-q), as described previously^15^. Because single SpsP neurons innervate the entire left or right side of the protocerebral bridge (Extended Data Fig. 3e, f (SpsP row), p-q), when imaging them we treated the entire left bridge as one ROI and the entire right bridge as another. When imaging PFNs or LNO1s in the noduli (Extended Data Figs. 3b-c, i, l-m), we treated the entire left nodulus as one ROI and the entire right nodulus as another.

For ellipsoid body imaging (Figs. 1, 3o, 4 and Extended Data Figs. 1, 6, 7), we defined ROIs by first outlining the region of each z-slice that corresponded to the ellipsoid body. We then radially subdivided the ellipsoid body into 16 equal wedges radiating from a manually defined center, as describe previously^12^. For fan-shaped body imaging (Figs. 1, 3o, 4 and Extended Data Figs. 1, 6, 7, 8f-g, l-m, r-s), we defined ROIs by first outlining the region in each z-slice that corresponded to the fan-shaped body. We then defined two boundary lines delineating the left and right edges of the fan-shaped body. When these two edge lines were extended down, they met at an intersection point beneath the fan-shaped body. We subdivided the angle generated by thus intersecting the two fan-shaped body edges––which corresponds to the overall angular width of the fan-shaped body region––into 16, equally spaced, angular subdivisions radiating from the intersection point. We assigned pixels to one of the sixteen fan-shaped body columns based on the pixel needing to (1) reside in the overall fan-shaped region and (2) reside in the radiating angular region associated with the column of interest.

### Calculating fluorescence intensities

We used ROIs, defined above, as the unit for calculating fluorescent intensities (see above). If pixels from multiple z-planes corresponded to the same ROI (e.g., the same column in the fan-shaped body), as defined above, then we grouped pixels from the multiple z-planes together for generating single fluorescence signal for that ROI. For each ROI, we calculated the mean pixel value at each time point and then used three different methods for normalization. We call the first method *ΔF/F*_*0*_ (Figs. 2d-n, 3m and Extended Data Figs. 2, 3i PFN_d_s and PFN_d_s in the bridge, j, l, n, p, 4, 8), where F_0_ was defined as the mean of the lowest 5% of raw fluorescence values in a given ROI over time and ΔF was defined as F – F_0_. We call the second method *normalized ΔF/F*_*0*_ (Figs. 1, 2c, 3o, 4, and Extended Data Figs. 1, 6, 7), which uses this equation: (F – F_max_)/(F_max_-F_0_), where F_0_ was still the mean of the lowest 5% of raw fluorescence values in a given ROI over time and F_max_ was defined as the mean of the top 3% of raw values in a given ROI over time. This metric normalizes the fluorescence intensity of each glomerulus, wedge or column ROI to its own minimum and maximum and makes the assumption that each column, wedge or glomerulus has the same dynamic range as the others in the structure, with intensity differences arising from technical variation in the expression of indicator or from the number of cells expressing indicator within a column or wedge. We used this method to estimate the phase of heading/traveling signals where it seemed reasonable to make the above assumption for accurately estimating the phase of a bolus in a structure. We call the third method *z-score normalized ΔF/F*_*0*_ (Extended Data Figs. 3b-h, i signals in the noduli and SpsPs, k, m, o, q) where we show how many standard deviations each time point’s signal is away from the mean. We calculated the signal as *ΔF/F*_*0*_ and then we z-normalized the signal. We used this method to estimate the asymmetry of neural responses to optic flow in the bridge or noduli, where it seemed sensible to normalize the baseline asymmetry (when there are no visual stimuli) to zero. Importantly, none of the conclusions presented in this paper rely on the normalization method used for visualizing and analyzing the data.

### Calculating the phase of boluses and aligning phase across structures

To calculate the phase of the joint movement of the calcium boluses in the left and right protocerebral bridge, we first converted the raw bridge signal into a 16-to-18-point vector, with each glomerulus’ signal normalized as described above. Then, for each time point, we took a Fourier transform of this vector and used the phase at a period of 8 glomeruli to define the phase of the boluses, as previously described^15^. To calculate phase of the EPG bump in the ellipsoid body, we computed the population vector average of the 16-point activity vector, as previously described^12^. To calculate the phase of the PFR bolus in the fan-shaped body, we computed the population vector average like in the ellipsoid body, using the following mapping of fan-shaped body columns to ellipsoid body wedges. The leftmost column in the fan-shaped body corresponded to the wedge at the very bottom of the ellipsoid body, just to the left of the vertical bisecting line; the rightmost column in fan-shaped body corresponded to the wedge at the very bottom of the ellipsoid body, just to the right of the vertical bisecting line. We then numbered the fan-shaped body columns 1 to 16, from left to right, just like we numbered the ellipsoid body wedges clockwise around that structure^12^. This mapping is meant to match the expected mapping of signals from anatomy, described previously^33^, and as further discussed immediately below.

To align the EPG phase in the ellipsoid body with the PFR phase or the hΔB phase in the fan-shaped body, we used the approach just described (Figs. 1, 3, 4, and Extended Data Figs. 1, 6, 7). To align the EPG and PFN_d_ and PFN_v_ phase signals in the protocerebral bridge (Fig. 2c-e), we used the fact that these neuron populations commonly innervate 14/18 glomeruli in the protocerebral bridge, which allows for an obvious alignment anchor, as done previously^15^. To calculate the offset between the phase of neural boluses and the angular position of a cue (bright bar or dot) rotating in angular closed loop on our visual display, we computed the circular mean of the difference between the neural phase and the cue angle during the time points when the cue was visible to the fly. We used this difference to provide a constant (non-time-varying) offset to the neural phase signal such that the difference between the phase and cue angles was minimized across the whole measurement window of relevance. This approach is needed because of the past finding that phase signals in the central complex have variable offset angles to the angular position of cues in the external world across flies (and sometimes across time within a fly)^12,15^. To calculate the phase offset between neural bolus position and visual cue angle, we did not analyze time points when the fly was not flying in all of our flight experiments (Figs. 1-4, and Extended Data Figs. 2, 3, 4, 6-8) nor did we analyze time points when the fly was standing in walking experiments (Fig. 4c, d). For a fly to be detected as standing, the forward speed needed to be less than 2 mm/s, the sideslip speed less than 2 mm/s, and the turning speed less than 30 °/s. We also excluded the first 10 s of each period in Figure 1f-g to minimize the impact of a changing visual stimulus on the offset estimate.

### Comparing data acquired at different sampling rates or with a time lag

When comparing two-photon imaging data (collected at ∼5-10 Hz) and behavioral (flight turns or ball walking) data (collected at 50-100 Hz) for the same fly, we subsampled the behavioral data to the imaging frame rate by computing the mean of behavioral signals during the time window in which each imaging data point was collected (Figs. 1f, 4b-d and Extended Data Figs. 1f, 3j-q), as previously described^15^.

Although we collected both the EPG signal in the ellipsoid body and the PFR signal in the fan-shaped body at the same frame rate, the precise time points in which these two signals were sampled were slightly different because the piezo drive that moves the objective had to travel from the higher fan-shaped body z-levels to the lower ellipsoid body z-levels. Importantly, each z-slice in a such volumetric time series was associated with its own trigger time and we could use this fact to more accurately align the fan-shaped body and ellipsoid body phase signals to each other. Specifically, when comparing EPG and PFR bolus positions over time, we first created a common 10 Hz (100 ms interval) time base. We then assigned phase estimates from the two structures/cell-types to this common time base by linearly interpolating each time series (using its specific z-slice triggers) and we used these interpolated time points, on the common time base, for calculating the phase differences between EPGs and PFRs, or EPGs and hΔBs (Figs. 1h-o, 3o and Extended Data Figs. 1e, f, 6, 7). For the histograms and other analyses in Figures 1g, 4g, h, k, l, o, p, we simply subtracted the EPG phase and the PFR phase measured in each frame, without temporal interpolation. None of our conclusions are altered if we change the interpolation interval or do not interpolate.

### Backward walking analysis

We expressed CsChrimson in a group of lobula columnar neurons, LC16, whose activation with red light has been shown to induce flies to walk backward (Extended Data Fig. 1c-f, more details in ‘Optogenetic stimulation’)^47^. Consistent with previous studies in free walking flies^47^, we also observed variable backward walking behaviors mixed with sideward walking and turning in our tethered preparation. To test whether the PFR phase separates from the EPG phase when a fly walks backward, we analyzed optogenetic activation trials based on the following three criteria being met. First, the backward walking speed needed to be larger than 6 mm/s. Second, the duration of continuous backward walking (defined by backward walking speed being above 0.5 mm/s) needed to be longer than 1 s. Third, during the backward walking period, the sideward walking velocity needed to be biased toward one direction; the fraction of optogenetic trials in which the sideward velocity was clearly either positive or negative exceeded 80%. We included this third criterion so that we could split optogenetic trials into those where the PFR phase should have moved to the right and those in which it should have moved to the left in the fan-shaped body (Extended Data Fig. 1c-f).

### Phase nulling

To compute the time-averaged shape of the bolus in PFNs and EPGs, we followed previous methods^15^. In brief, we (1) computationally rotated each frame by the estimated phase of the bolus on that frame, such that the bolus peak was at the same location on all frames and then (2) averaged together the signal from all frames to get an averaged bolus, whose shape we could analyze via fits to sinusoids (Fig. 2 and Extended Data Figs. 2, 4, 8a). In this phase nulling process, we first interpolated the GCaMP signal from each frame to 1/10 of a glomerulus, column, or wedge with a cubic spline. We then shifted this interpolated signal by the phase angle calculated for that frame. In both the ellipsoid body and fan-shaped body, we performed a circular shift, such the signals wrapped around the edges of the fan-shaped body. In the protocerebral bridge, we performed this circular shift independently for the left and right bridge. For the protocerebral bridge data, we subsampled the spatially interpolated GCaMP signal back to a 16-glomerulus vector before plotting the data (Fig. 2 and Extended Data Figs. 2, 4, 8a) so as to more accurately reflect, in our averaged signals, what the actual signal in the brain looked like.

### Modeling

We constructed and analyzed an analytical model of the synaptic input to hΔB neurons to understand how the traveling direction determines the angular location of the peak hΔB activity across the fan-shaped body. Neurons in the model are labeled by an angle *θ*that indicates the glomerulus in the protocerebral bridge innervated by a particular neuron or, in the case of hΔB, its position in the fan-shaped body. In reality, this angle label takes discrete values (Extended Data Fig. 4b, j, k) but, to simplify the notation, we use a continuous label here.

The fly’s allocentric heading angle is denoted by *H*, and its allocentric traveling direction angle by *T*. The egocentric traveling angle is then *T*-*H*. Another relevant angle is the angular location of the bolus of EPG activity in either the ellipsoid body or the protocerebral bridge. All these angles are measured relative to an arbitrary axis and, for simplicity, we choose this axis so that the bump angle is −H, the negative of the heading angle. The minus sign reflects the fact that when the fly turns clockwise the bump rotates counterclockwise.

To make things less confusing with regard to this minus sign, we flipped the orientation of the horizontal axis in some of our figures.

The activity of each PFN class forms a sinusoidal pattern across the protocerebral bridge (Fig. 2d, e) and fan-shaped body. We are interested in constructing the PFN input to the hΔBs, so we model these patterns in the fan-shaped body:

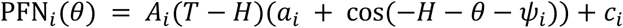

where *i* = 1, 2, 3, 4 refers to right-bridge PFN_d_, left-bridge PFN_d_, right-bridge PFN_v_, and left-bridge PFN_v_, and *a*_*i*_and *c*_*i*_are free parameters. *A*_*i*_(*T* − *H*)is the amplitude of the sinusoid for PFN *i* when the egocentric traveling angle is *T* − *H* (Fig. 2g, h, j, k). The angles *ψ*are the shifts in the PFN-to-hΔB projections (Fig. 3n). The above equation implies that the mean PFN activity across *θ* values is

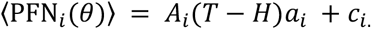

We fit the mean PFN activities by using the data points for *A*_*i*_(*T* − *H*)and adjusting the parameters *a*_*i*_ and *c*_*i*_(Extended Data Fig. 8n-q). The above equation also allows us to extract the amplitudes *A* from the means ⟨PEN(*θ*)⟩, a feature we took advantage of when using data obtained from the noduli. We fit the PFN amplitudes as shifted sinusoids,

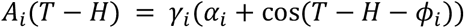

where *α*_*i*_, *γ*_*i*_ and *ϕ*_*i*_ are free parameters. The fits obtained from this formula are shown in Fig. 2g, h, j, k for one speed, and in Extended Data Fig. 8b-e for all four speeds tested.

The input to the hΔBs is given by the sum of PFN activities weighted by factors *g* that reflect the strength of their connections to the hΔB,

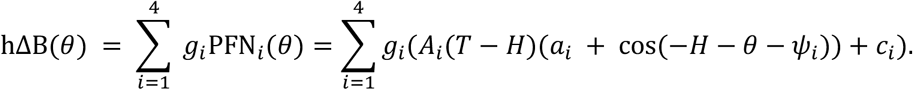

For reasons provided below, we choose the *g* so that *g*_*i*_*γ*_*i*_*α*_*i*_ = *C* for all *i*, where C is a single constant that scales the hΔB response. The traveling direction angle encoded by the model (Fig. 3o) is the value of *θ*that maximizes the above expression for a given value of *T* − *H*. To generate the results in Fig. 3o, we used the data points for the amplitudes *A*_*i*_(*T* − *H*). Note that the maximizing *θ*value does not depend on the parameters *a*_*i*_, *c*_*i*_or *C*, which means that the model results in Fig. 3o do not involve any additional parameters beyond what we extracted from the PFN and EM data.

We also modeled the input to the PFR neurons as the sum of the activities of their inputs from hΔB, PFN_d_ and EPG neurons. The EPG inputs to the PFRs appear to be weak, and we consistently found that the best fits to the data were obtained with the strength of the EPG-to-PFR connection set to zero. Therefore, we leave the EPG input out of this discussion. Using the remaining inputs, we have

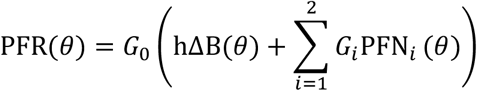

where *G*_0_ is an overall scale factor, and *G*_1_ and *G*_2_ are relative input-strength factors. We used the data points for the responses of the hΔB and PFN inputs in the above equation to generate the fit in Extended Data Figure 6e. Because *G*_0_ does not affect this result and we imposed the constraint *G*_1_ = *G*_2_, this fit involves one free parameter.

We now provide a complete analysis of the model outlined above. To simplify the model, we note that the values of *α*_*i*_ for all *i* are close to each other, so we replace all four with a single parameter 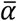. We assume that all four PFNs contribute equally to the hΔB activity (although this assumption is not essential) so, as above, we set *g*_*i*_*γ*_*i*_*α*_*i*_ = *C* for all *i* but, in this case, the condition is 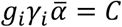. As a result *g*_*i*_*γ*_*i*_ = *D* for all i, where 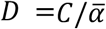 is another constant. Under these assumptions

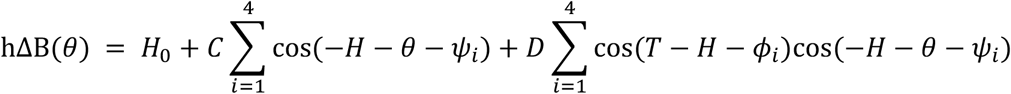

where *H*_0_ denotes all the terms that do not depend on *θ*. We do not need to consider these terms in our analysis because we are interested in deriving the profile of the hΔB input across the fan-shaped body, that is the activity as a function of *θ*, not the mean level of this activity.

The angles *ψ* are spaced from each other by differences close to 90° (Fig. 3n), suggesting that the 4 PFNs are acting as an orthogonal system. In this case, for vectors to be represented faithfully, the angles *ψ*, reflecting the anatomy of the PFN projections, and the angles *φ*, that reflect the physiology of the PFN responses, should be equal to each other, that is, *ψ*_*i*_ = *ϕ*_*i*_ for all *i*. The data (Fig. 3n, m) suggest that these common angles are 45°, −45°, −135° and 135°. With these angular values, the middle term on the right side of the above equation is zero, and summing the third term gives

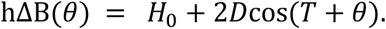

This means that the hΔB input is maximal at *θ*_max_ = −*T*, indicating that the angle of the hΔB bolus is aligned with the traveling direction angle. The minus sign implies that the hΔBs encode minus the traveling-direction angle, −*T*, just as the EPGs encode minus the heading angle, −*H*.

An interesting feature of the sinusoidal representation of 2-D vectors is that it nicely accommodates non-orthogonal coordinate systems. Our estimate of the angles *ψ*involves a number of assumptions and, as mentioned in the Discussion, it is possible that these angles can even change in different contexts (such as walking vs. flying). If these angles are not separated by 90°, this implies a non-orthogonal coordinate system. In this case, the two sets of angles, *ψ* and *ϕ*, are not equal, but there is still a definite relationship between them. Assuming a noise-robust solution, this relationship can be derived by performing a pseudo-inverse operation on a matrix of unit vectors oriented at the angles *ψ*to derive *ϕ*, or at the angles *ϕ* to derive *ψ*. In addition, the coupling strengths *g* need to be adjusted to accommodate the lengths of the vectors arising from this computation.

## Data and code availability

All data and analysis code are available upon request.

---

**Extended Data Figure 1.**
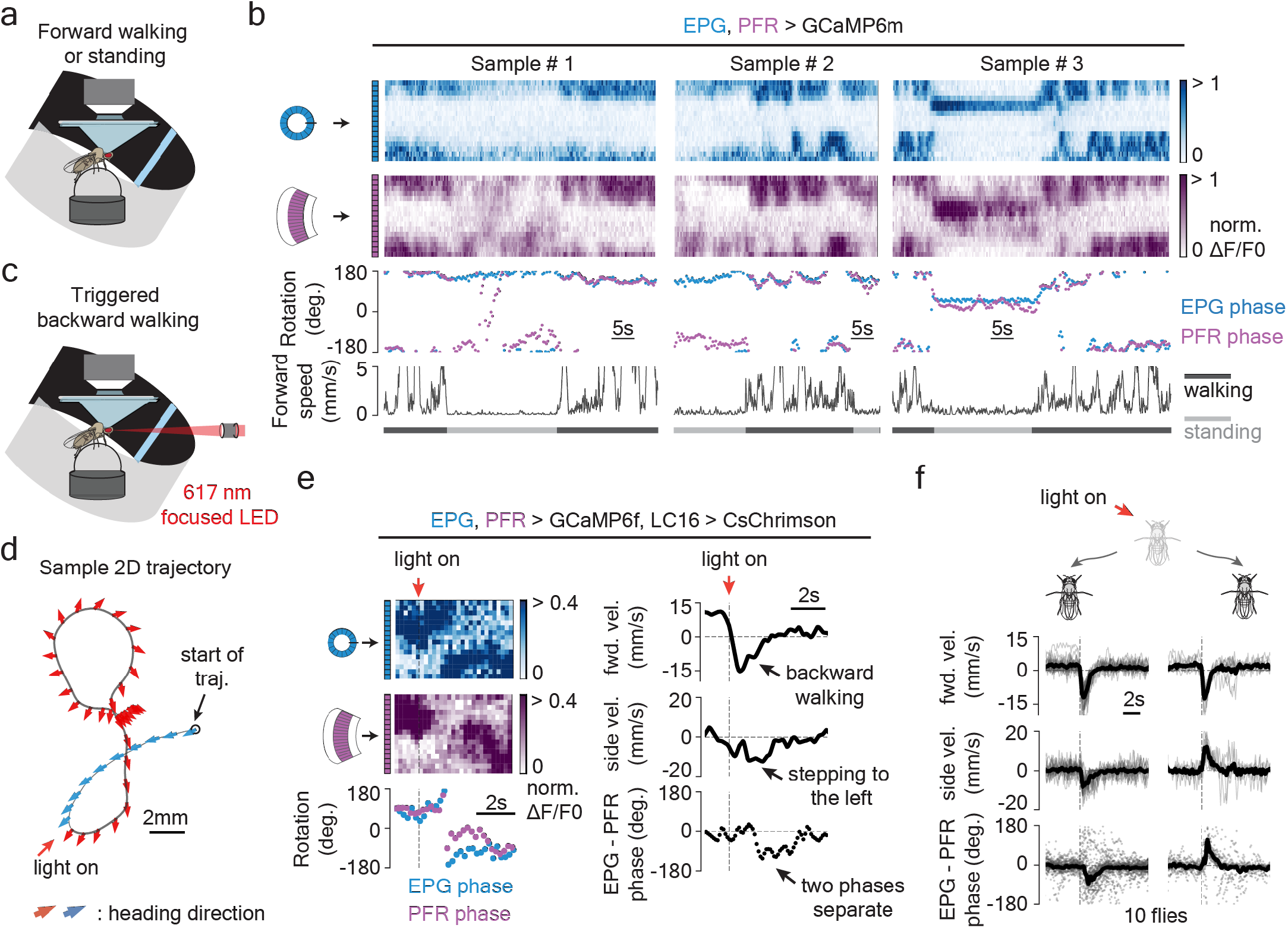
The PFR bolus position drifts in standing flies, and it deviates from the EPG bolus position when flies walk backward in a manner consistent with the PFR phase signaling the fly’s traveling direction during walking. **a**, Tethered, walking, [Ca^2+^]-imaging setup with a bright blue bar that rotates in closed loop with the fly’s turns. **b**, Sample time series of simultaneously imaged EPG and PFR boluses in a tethered, walking fly. Top two traces show [Ca^2+^] signals. Third trace shows the phase estimates of the two boluses. Bottom trace shows the forward speed of the fly. **c**, Tethered-walking setup where we used a 617 nm LED focused on the center of the fly’s head to optogenetically trigger backward walking via activation of LC16 visual neurons expressing CsChrimson^47^ (Methods). **d**, An example 2D trajectory of optogenetically triggered backward walking. An arrow is shown every ∼0.1 seconds. Red arrows indicate backward walking during the red-light pulse; blue arrows indicate the 1.2 s before the red light turned on. **e**, Left, time series of EPG (blue) and PFR (purple) boluses and phase-estimates from the trajectory in panel d. Right, time series of forward velocity, sideslip velocity and the difference between the PFR and EPG phase in the trajectory shown in panel d. The ΔF/F heatmap range is more compressed here than in other plots because the PFR signal strength typically dips when the fly initiates backward walking (a phenomenon whose mechanism we have not yet explored). Nevertheless, clear moments where the PFR phase separates from the EPG phase are evident, even after the PFR signal strength has recovered, in this sample trace (and in others). **f**, Time series of the mean forward velocity, mean sideslip velocity and the circular mean of the difference between the PFR and EPG phase during backward walking, grouped by optogenetic trials in which the fly walked to the back left (left panel) or to the back right (right panel). The sign of PFR-EPG phase deviations seen here, in walking, are consistent with the signs observed in flight, for the same directions of backward-left and backward-right travel. Thin lines and gray dots: individual trials. Thick line and black dot: population mean (circular mean for bottom row).

**Extended Data Figure 2.**
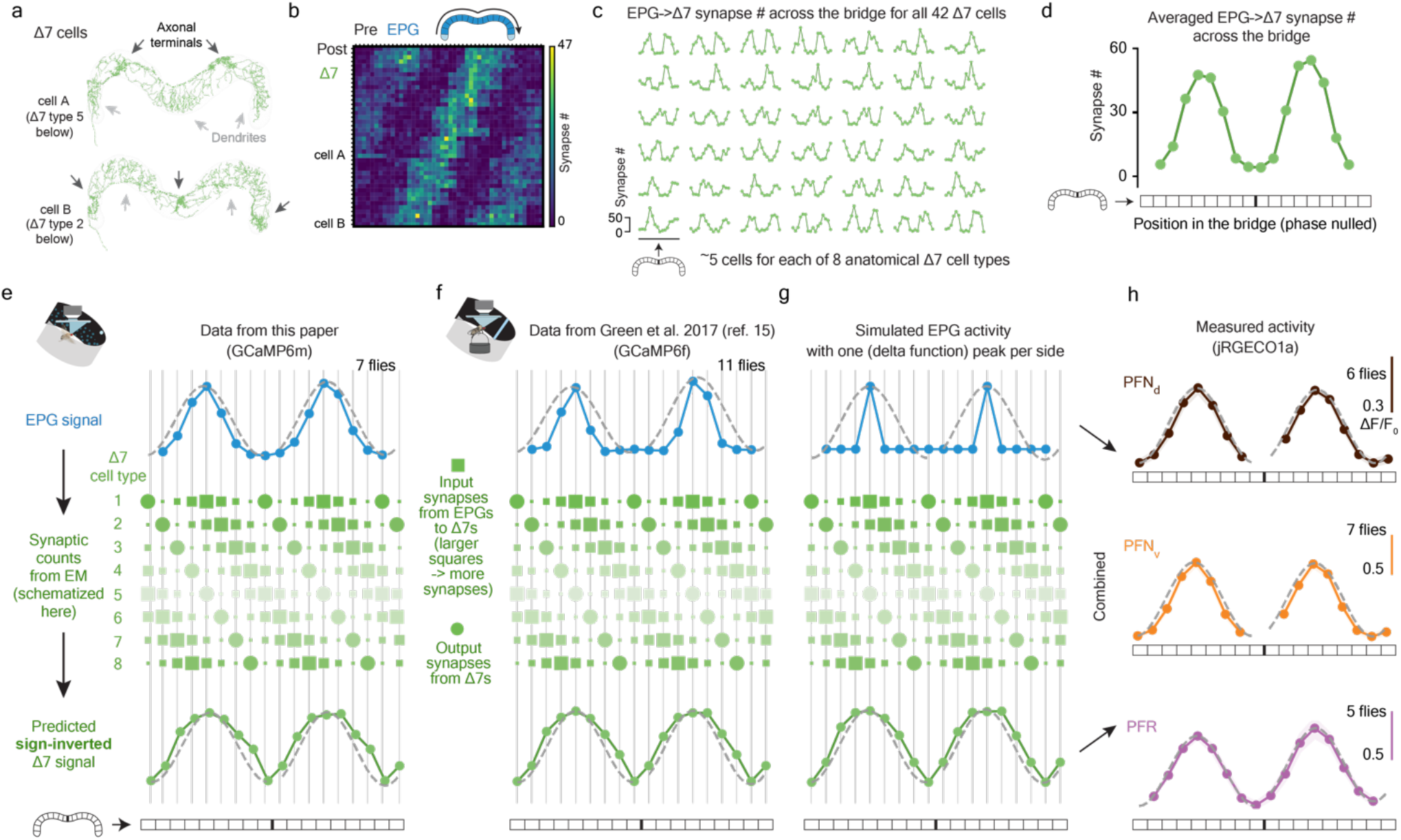
Δ7 cells are poised to help create sinusoidally shaped activity boluses in PFN_d_s, PFN_v_s and PFRs in the protocerebral bridge. Connectivity data are based on those in neuPrint^38^, hemibrain:v1.1. **a**, Two Δ7 cells from neuPrint reveal a graded increase and decrease in dendritic density across the bridge. **b**, Synapse-number matrix for detected synapses from EPG cells to Δ7 cells in the protocerebral bridge. Each row represents one Δ7 cell. **c**, Same data as in panel b, but plotting each Δ7 cell separately. **d**, Phase-nulled EPG-to-Δ7 synapse # across the glomeruli of the bridge, averaged across all 42 Δ7 cells, based on the data in panel c. The anatomical input strength from EPGs to Δ7s is sinusoidally modulated across the bridge. **e**, Transforming the EPG activity pattern across the bridge (blue) into a predicted Δ7 activity pattern (green, bottom row) based on the synaptic density profile in panel c (schematized in the middle). We first calculated the dot product between the EPG activity vector and each Δ7 cell’s EPG-to-Δ7 synapse-number vector (panel c). Then, for each glomerulus, we averaged the dot-product-output for all of the Δ7s that have axonal terminals in that glomerulus, thus creating the predicted activity value for that glomerulus. (The size of each green square here schematizes the # of synapses from EPGs to the Δ7 cell of that type in that column; the intensity of each Δ7 row indicates the expected output strength of each Δ7 cell type, after being driven by the EPG signal above.) We plot the inverted, predicted activity output from Δ7s in the bottom row (green) because Δ7s are glutamatergic^48^ and glutametergic neurons in the *Drosophila* central nervous system typically inhibit their postsynaptic targets (via Glu-Cl channels). After inverting the Δ7 activity one can then imagine simply averaging the Δ7 predicted-activity row with the EPG activity––with some relative weighting for the Δ7 and EPG curves––to generate the net drive to the many downstream neurons that receive both EPG and Δ7 input^38^, like PFNs. Note that the EPG activity boluses are slightly narrower than the sinusoidal fits whereas the Δ7 activity boluses are slightly wider than the sinusoidal fits. **f**, Same as panel e, but using the phase-nulled, averaged EPG GCaMP activity pattern from a previous study^15^. Note although the EPG bolus is narrower in these data from walking flies than in panel e from flying flies, the shape of the predicted Δ7 output remains similar. **g**, Same as panel e, but starting with (imagined) EPG activity where there is only one active glomerulus on each side of the bridge. Note that the shape of the predicted Δ7 output remains similar to that in panels e-f. **h**, Measured, phase-nulled activity profiles from PFN_d_s, PFN_v_s and PFRs. All three activity patterns conform well to their sinusoidal fits (gray dashed lines). We hypothesize that the sinusoidal activity patterns in bridge columnar cells like PFN_d_s, PFN_v_s, PFRs arises from the combined impact of EPG and Δ7 input. In other words, we posit that Δ7s “sinusoidalize” the EPG bumps in the bridge –– that is, they function to broaden and smoothen the EPG input to the bridge, to create two sinusoidally shaped boluses in their recipient cells, with these boluses often functioning as explicit, 2D vector signals in the fan-shaped body.

**Extended Data Figure 3.**
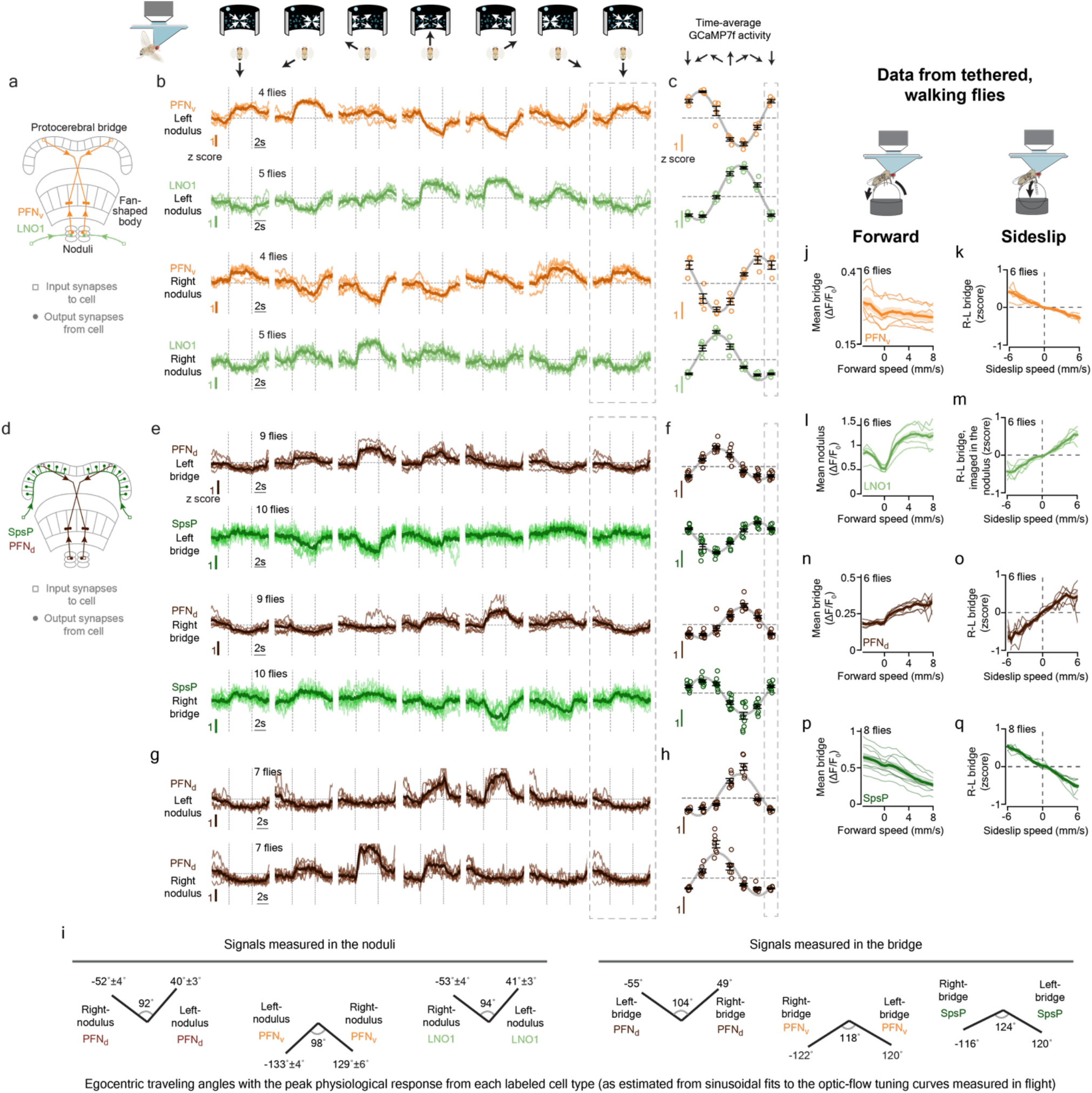
LNO1 and SpsP cells have [Ca^2+^] responses that are strongly tuned to the fly’s egocentric translation direction––in both walking and flying flies––with responses suggesting that these cells provide sign-inverting, inhibitory input to PFN_v_ and PFN_d_ cells, respectively. **a**, LNO1s are a class of cells (two total neurons per side, four per brain) that receive extensive synaptic input outside the central complex and provide extensive synaptic input to PFN_v_s in the noduli^38^. **b**, Mean GCaMP signals in PFN_v_s and LNO1s in the nodulus as a function of the simulated traveling direction of the fly (via open loop optic flow). Dotted rectangle indicates a repeated-data column, in this panel and throughout. **c**, Single-fly (colored circles) and population means ± s.e.m. (black bars) of the average signal in the final 2.5 s of the optic flow epoch. **d**, Each SpsP cell (two total neurons per side, four per brain) receives extensive synaptic input outside the central complex and provides extensive synaptic input to PFN_d_s on one side of the protocerebral bridge^38^. **e**, Same as panel b, but mean GCaMP signals in PFN_d_s and SpsPs in the bridge as a function of the simulated traveling direction of the fly (via open-loop optic flow). A closed-loop bright dot was not present on the LED display when collecting the PFN_d_ data. **f**, Same as panel c, but averaging the bridge signal in panel e. **g**, Same as panel b, but analyzing the PFN_d_ signal in the noduli. A closed-loop bright dot was not present on the LED display. **h**, Same as panel c, but averaging the nodulus signal in panel g. **i**, The optic-flow-simulated egocentric traveling angle at which the activity of each cell type is strongest is depicted with a line at the associated angle. Note that the left-vs.-right angular differences measured in the noduli are smaller, and closer to 90°, than the left-vs.-right angular differences measured in the bridge. **j**, Data collected from tethered flies walking on a floating ball in complete darkness are shown in this panel and all subsequent panels in this figure. Mean PFN_v_ GCaMP signals in the bridge as a function of the fly’s forward speed. **k**, Right-minus-left PFN_v_ GCaMP signals in the bridge as a function of the fly’s sideslip speed. **l-m**, Same as panel j and k, but analyzing LNO1 signals in the nodulus. **n-o**, Same as panel j and k, but analyzing PFN_d_ signals in the bridge. **p-q**, Same as panel j and k, but analyzing SpsP signals in the bridge. In panel b, e, g, j-q, thin lines represent single-fly means and thick lines represent population means. Note that PFN_v_s and LNO1s have sign-inverted responses, and that PFN_d_s and SpsPs have sign-inverted responses. The response signs to optic-flow simulating the fly’s body translating forward and leftward (rightward) in flight are the same as the signs of responses to the fly walking forward and side-slipping leftward (rightward) when walking. Thus, these data are consistent with all these neurons being sensitive to the fly’s egocentric translation direction, as assessed via optic flow (dominantly) in flight, and via proprioception or efference-copy (dominantly) in walking.

**Extended Data Figure 4.**
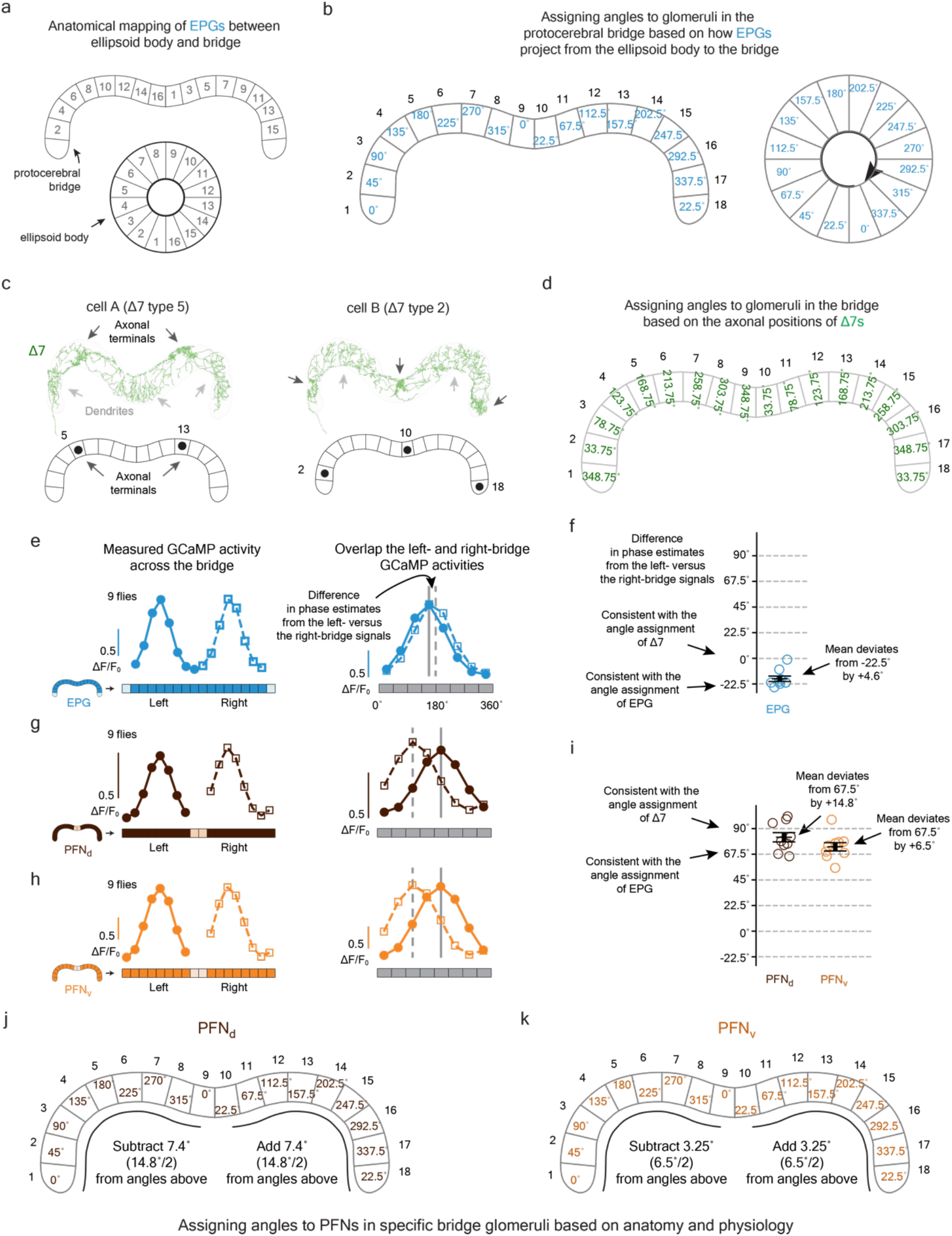
Multiple, functionally relevant ways of indexing angles across the protocerebral bridge. Connectivity data and cell-type names are based on those in neuPrint^38^, hemibrain:v1.1. **a**, The previously described mapping between EPG dendritic locations in the ellipsoid body and axonal-terminal locations in the bridge^33^. Numbers ordered based on the location of each EPG cell in the ellipsoid body. **b**, EPG cells divide the ellipsoid body into 16 wedges, each 22.5° wide. Each glomerulus in the bridge inherits its angle, in our analysis here, based on the EPG projection pattern shown in panel a. The angles of the outer two bridge glomeruli––which do not receive standard EPG input, but only EPGtile input^38^––were inferred to have angles equal to the middle two glomeruli (0° and 22.5°, respectively) based on how other cell types innervate the bridge, as discussed in past work^15^. Glomeruli are numbered 1 to 18 from left to right, to aid the comparisons made below. **c**, Two Δ7 cells from neuPrint (and past work^33^) reveal that the axonal terminals of each Δ7 cell are 8-glomeruli apart (#5→#13 for cell A and #2→#10→#18 for cell B). This anatomy argues that any two glomeruli 8 apart, such as #5 and #13, will experience Δ7 output of equal strength. Compelling physiological evidence for this statement is available in the [Ca^2+^] signals of the PEN2 (equivalently, PEN_b) columnar cell class in the bridge, which is a strong anatomical recipient of Δ7 synapses^38^ and shows [Ca^2+^] activity across the bridge––clearly dissociable from the activity in EPGs––with consistently equal signal strength at glomeruli spaced 8 apart, perfectly following the Δ7 anatomical prediction (orange trace in Fig. 3d and data points in Extended Data Fig. 2i from ref. 15). Note that in the EPG indexing, shown in panel b, glomeruli #5 and #13, as examples, have angular indices that are not identical, but differ by 22.5°. **d**, Angles assigned to each bridge glomerulus based on the Δ7 axonal anatomy. Because the Δ7 output anatomy requires that any two glomeruli 8 apart, across the whole bridge, have the same angular index assignment, this results in a situation where all neighboring glomeruli have angular assignments that are separated by 45°. Note that almost all neighboring glomeruli are separated by 45° in the EPG mapping as well, except that, critically, in the EPG mapping the middle two glomeruli are separated by only 22.5°. This discontinuity is not evident in the Δ7 output. To create an angular indexing of the bridge for Δ7s that accommodates the anatomical constraints just described––i.e., one that incorporates an additional 22.5° in the bridge representation of angular space and thus “erases” the EPG discontinuity––we shifted the angular index for each glomerulus on the left bridge leftward by 11.25° relative to the EPG indexing and we shifted the angular index for each glomerulus on the right bridge rightward by 11.25° relative to the EPG indexing. **e**, The EPG indexing in panels a-b predicts that EPG activity in the left bridge (#2→#9) will be left-shifted by 22.5° compared to EPG activity in the right bridge (#10→#17). Indeed, when we overlapped the left- and right-bridge EPG signals we found the two curves are detectably offset from each other. **f**, To quantify the data from panel e, for each imaging frame in which the fly was flying, we calculated the phase of the EPG bump in the left and right bridge separately (via a population-vector average) and took the difference of these two angles. We then averaged this angular difference across all analyzed frames for the same fly. For EPGs, this angular difference should be –22.5° if it follows the EPG indexing in panel b and it should be 0° if the activity follows the Δ7 indexing in panel d. Across a population of 9 flies, we found the angular difference is close to –22.5°, but shifted toward 0° by 4.6°, consistent with the fact that the EPG signal itself receives strong anatomical input from the Δ7s and thus could be modulated in its shape to follow the Δ7 indexing, in principle^38^. It seems that the Δ7 feedback to EPGs reshapes its signal, but incompletely. **g-h**, Same as panel e, but analyzing the PFN_d_ and PFN_v_ activity in the bridge. Because PFNs only innervate the outer 8 glomeruli in each side of the bridge (unlike EPGs, which innervate the inner 8), we compared glomeruli #1→#8 in the left bridge overlapped with glomeruli #11→#18 in the right bridge here (the middle two glomeruli contain no signal for PFNs). **i**, Same as panel f, but analyzing the PFN_d_ and PFN_v_ activity in the bridge. Note that because PFN_d_s and PFN_v_s innervate (and thus we can only analyze) the outer 8 glomeruli of the bridge, the angular difference in phase estimates between the left- and right-bridge activity should be +67.5° if it follows the EPG indexing (panel b) and +90° if it follows the Δ7 indexing (panel d). We found that the average angular difference in both PFN_d_s and PFN_v_s is intermediate between +67.5° and +90°, consistent with PFNs receiving functional inputs from both EPGs and Δ7s. We use the angular offsets measured in this panel as the basis for slightly adjusting the PFN_d_ and PFN_v_ angular indices in the bridge to an intermediate value between the EPG and Δ7 indexing options, described above. We believe that this approach represents the most careful way to combine the known anatomy and physiology to determine the azimuthal angle that each PFN cell signals with its activity in driving the hΔB neurites in the fan-shaped body, which we analyze in the next figure. **j**, Angles assigned to each bridge glomerulus for PFN_d_s, based on the EPG indices from panel b and the physiologically determined adjustment required, based on the measurements in panel i. **k**, Same as panel j, but for PFN_v_s.

**Extended Data Figure 5.**
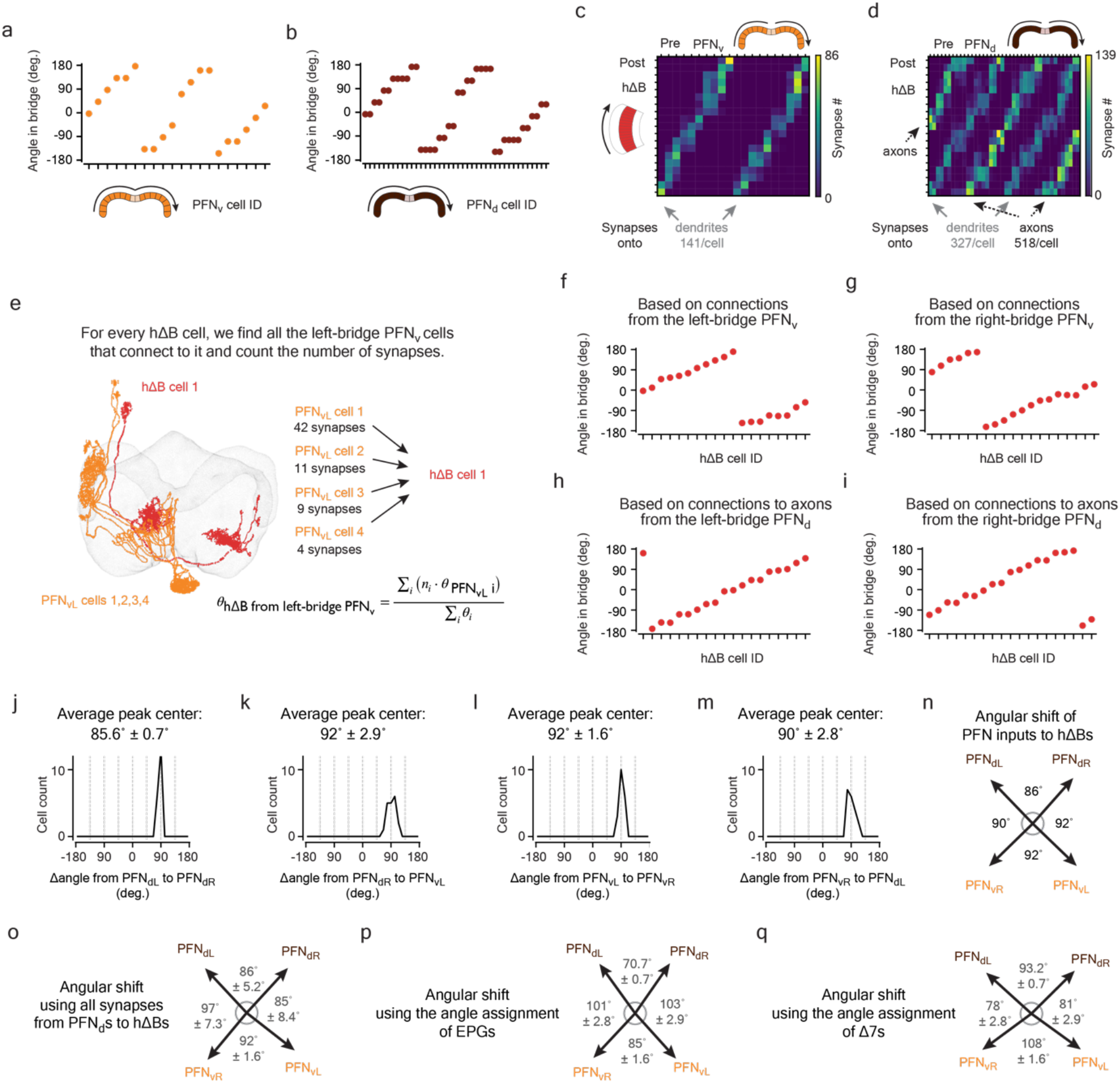
Computing the angular shift implemented by the PFN-to-hΔB connections. Connectivity data and cell-type names are based on those in neuPrint^38^, hemibrain:v1.1. **a**, The anatomical angle of each PFN_v_ cell is indicated based on which glomerulus it inervates in the protocerebral bridge, using the indexing described in Extended Data Figure 4k. **b**, Same as panel a, but for PFN_d_s, using the indexing described in Extended Data Figure 4j. **c**, Synapse-number matrix for detected synapses from PFN_v_ cells to hΔB cells in the fan-shaped body. Note that the two stripes in the heatmap represent PFN_v_s synapsing onto the dendritic regions of hΔBs. **d**, Same as panel c, but for synapses from PFN_d_ cells to hΔB cells. Note that two of the five stripes in the heatmap represent PFN_d_s synapsing onto the dendritic regions of hΔBs, whereas the other, brighter, three stripes represent PFN_d_s synapsing onto the axons of hΔBs. The average # of synapses that each hΔB compartment (axon vs. dendrite) receives from PFN cells is indicated on the bottom. **e**, Because hΔBs are postsynaptic to both PFN_v_s and PFN_d_s that project to the fan-shaped body from both sides of the bridge (panel c-d), each hΔB cell can be assigned an anatomical angle in four potential ways. To calculate the angle for an hΔB cell through its connection with the left-bridge PFN_v_s, for example, we averaged the anatomical angles of all the left-bridge PFN_v_s that connect to the hΔB cell in question, weighted by the number of synapses from that PFN_v_ cell to the hΔB cell. **f**, The anatomical angle of each hΔB cell calculated based on its monosynaptic inputs from left-bridge PFN_v_s using the method described in panel e and data in panel c. **g**, Same as panel f, but calculations were made with right-bridge PFN_v_ inputs to hΔBs. **h**, Same as panel f, but calculations were made with left-bridge PFN_d_ inputs to hΔBs, using only the synapses formed on the axonal terminals of hΔBs. (We test the impact of this assumption––of complete functional dominance of PFN_d_ axonal synapses to hΔBs––below.) **i**, Same as panel f, but calculations were made with right-bridge PFN_d_ inputs to hΔBs, using only axonal synapses. **j**, For each hΔB cell, we calculated the angular difference between the mean left-bridge PFN_d_ input and the mean right-bridge PFN_d_ inputs (i.e., the difference between data points in panels h and i) and we plot a histogram of those values. **k-m**, same as j for the cell types indicated. **n**, The anatomically predicted angles for the coordinate axes of the four PFN vectors, as projected to the fan-shaped body and interpreted by hΔB axons and dendrites, calculated by averaging the histogram values in panels j-m, respectively. **o**, Same as panel n, but including all synapses from PFN_d_s to hΔBs, not just the axonal ones as used above. We weigh dendritic and axonal synapses by PFN_d_s to hΔBs equally in the panel-e calculation. Note that the angles between four coordinate-frame axes do not change very much when also including the dendritic synapses from PFN_d_s to hΔBs, likely because they are less numerous than the axonal ones and the impact of the dendritic angles also seem to cancel out in their net effect (compare panels o and n). **p**, Same as panel n, but using the EPG indexing from Extended Data Figure 4 instead of the adjusted PFN_v_ and PFN_d_ indexing. Note that the EPG indexing makes the front angle between the left- and right-bridge PFN_d_ axes smaller. The same is true for the back angle between the left- and right-bridge PFN_v_ axes. **q**, Same as panel n, but using the Δ7 indexing from Extended Data Figure 4 instead of the PFN_v_ and PFN_d_ indexing. Note that the Δ7 indexing makes the front and back angles broader than 90°, when used in isolation. This analysis suggests that EPG and Δ7 inputs to PFNs are perfectly weighted to create axes that are orthogonal in our experiments in flying flies and also raise the possibility that orthogonality of this 4-vector system can be dynamically modulated via changing the weights of EPG and Δ7 inputs to PFNs (see Discussion).

**Extended Data Figure 6.**
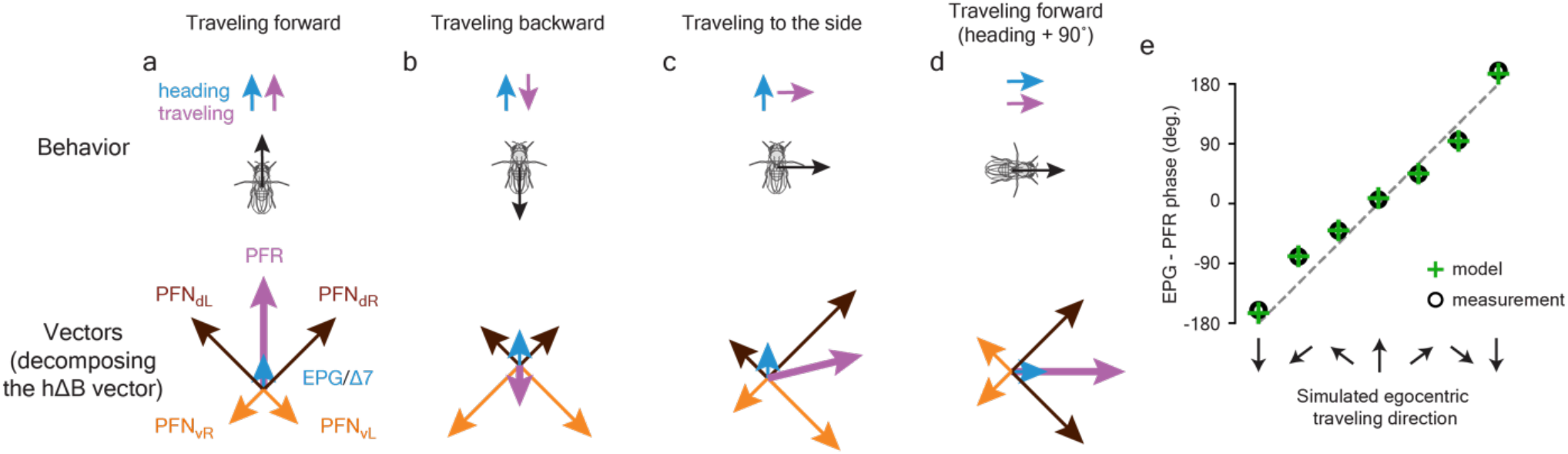
A five-vector model for building the PFR traveling-direction signal. PFRs receive hΔB input and thus they can be modeled to receive PFN_v_ and PFN_d_ vector input, disynaptically, through the hΔB route. PFRs also receive direct PFN_d_ monosynaptic input in the fan-shaped body and direct EPG/Δ7 monosynaptic input in the bridge. As such, one can model PFR cells as receiving five vector inputs: two PFN_v_ vectors whose magnitude is regulated by the PFN_v_→hΔB→PFR pathway exclusively, two PFN_d_ vectors whose magnitude is regulated by the PFN_d_→hΔB→PFR pathway augmented by additional, direct, monosynaptic inputs from PFN_d_s to PFRs, and one EPG/Δ7 vector input that arrives to the PFRs in the bridge. The weight of the EPG/Δ7 in our models always turns out to be small and thus that vector is depicted with a short blue arrow of unchanging length in the diagrams. **a**,When a fly travels forward, both PFN_d_ vectors are long and both PFN_v_ vectors are short, leading the sum of the five vectors, i.e., the PFR (purple) vector, to point forward (i.e., in the fly’s heading direction). **b**, When a fly travels backward, both PFN_d_ vectors are short and both PFN_v_ vectors are long, leading the sum, i.e., the PFR (purple) vector, to point backward. **c**, When a fly travels rightward, the right-bridge PFN_d_ vector and the left-bridge PFN_v_ vector are longer than their counterparts on the opposite side of the bridge, leading the sum, i.e., the PFR (purple) vector, to point rightward. Because PFRs receive––in addition to their hΔB inputs––direct, EPG/Δ7 input in the bridge and PFN_d_ input in the fan-shaped body, these additional inputs (if functionally relevant) will tend to bias the PFR traveling direction signal to the front. Indeed, in fitting the PFR data, our model gave these inputs (particularly the additional PFN_d_ input) functional weight because, empirically, we observed that the PFR bolus is indeed commonly skewed to the frontal direction (Figure 1i; PFR bolus position is consistently biased toward the horizontal zero line in cases where presented optic flow that simulated flies to be traveling sideways, ±60° and ±120°). While we do not yet functionally understand why PFRs incorporate a frontal skew to their traveling-direction signal, this skew is why the purple, PFR, summed vector points slightly upward here. **d**, Same as panel a, but after the fly has turned clockwise by 90°, which shifts the whole reference frame by 90° because both the PFN_d_ and PFN_v_ activites are linked to the EPG bolus, which rotates when the fly turns. **e**, There is a good correspondence between the measured deviation of the PFR phase from the EPG phase (as a function of the optic-flow direction presented) in Figure 1i (circles) and the deviation predicted by our formal model (plus signs). See Methods for details.

**Extended Data Figure 7.**
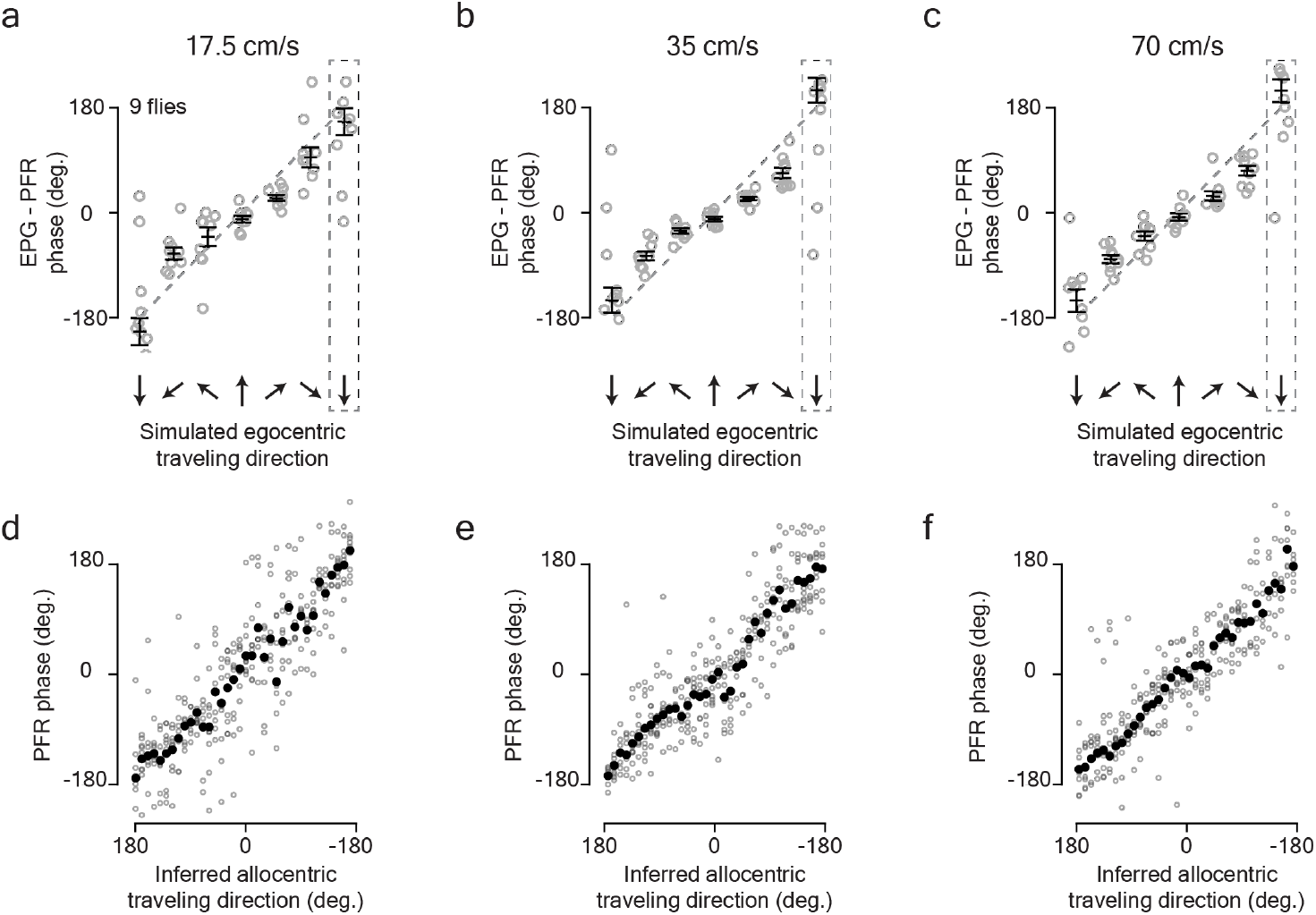
PFR cells signal *Drosophila*’s traveling direction at three different optic-flow speeds. **a**, Phase difference between EPGs and PFRs for different simulated traveling directions of the fly. The traveling direction was simulated with ventral/lateral optic flow, using a slow optic-flow speed (half the speed in Figure 1i). (See Methods for how we calculate the optic flow speed.) Circular means were calculated in the last 2.5 s optic flow presentation. Gray: individual fly (circular) means. Black: population (circular) mean and s.e.m. **b**, Same as panel a, but for a medium optic-flow speed (same speed as in Figure 1i). **c**, Same as panel a, but for a fast optic-flow speed (double the speed in Figure 1i). In panels a-c we note that the PFR-EPG phase difference is slightly above the unity line for optic-flow simulating rightward travel and slightly below unity for leftward travel. These deviations from linearity were not evident in the hΔB-EPG phase measurements (Figure 1n) and thus they are likely explicable by unique inputs to PFRs by comparison to hΔBs. Specifically, PFRs receive a direct EPG/Δ7 input in the bridge that hΔBs do not receive, as well as direct, monosynaptic PFN_d_ inputs that bypass the hΔBs, and are likely to make the impact of the PFN_d_ vectors stronger in PFRs over hΔBs. Both these additional inputs will tend to push the traveling-direction bolus to the forward direction, which is what the slight deviations from linearity in the above data reflect (see Extended Data Fig. 6 for further discussion). **d**, PFR phase as a function of the inferred allocentric traveling direction (see Main Text) in the context of a slow optic-flow speed. Gray: individual fly means. Black: population mean. Data were analyzed in the same way as in Figure 1i, j. **e**, Same as panel d, but for a medium optic flow speed. **f**, Same as panel d, but for a fast optic flow speed.

**Extended Data Figure 8.**
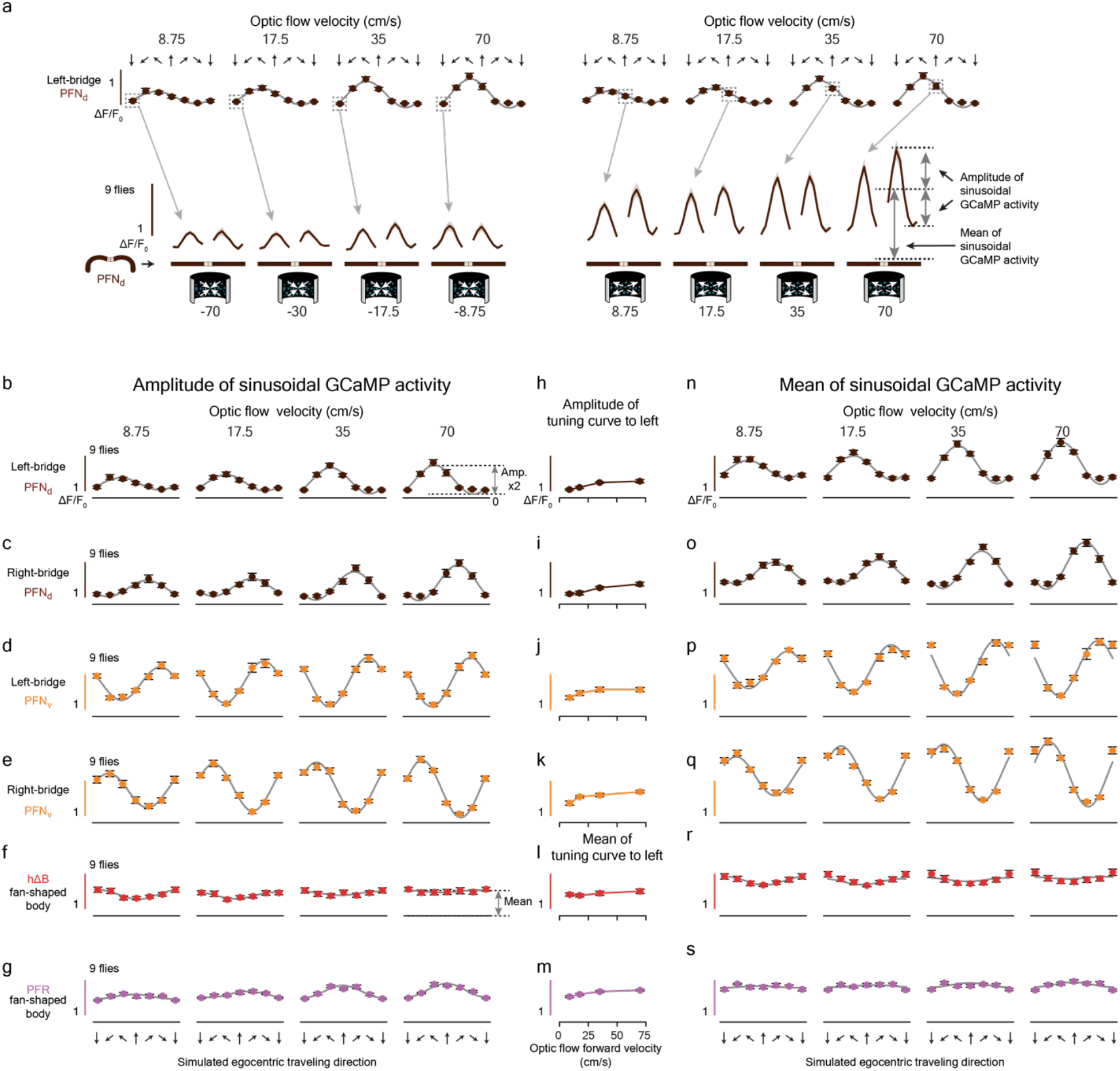
Response-tuning in PFN, PFR and hΔB neurons to the translation speed indicated by our optic-flow stimuli. **a**, Bottom row: phase-nulled PFN_d_ GCaMP activity across the bridge, averaged in the final 2.5 s of the optic flow epoch. We show responses to optic flow simulating traveling backward at four different speeds (left) and responses to optic flow simulating forward travel at four different speeds (right). The mapping between bridge [Ca^2+^] signals and data points in the plots in subsequent panels is indicated (arrows) for a few example points, using measurements from left-bridge PFN_d_s, as an example. How we calculate the mean and amplitude of each bolus is schematized. **b**, The population-averaged amplitude of the phase-nulled left-bridge PFN_d_ [Ca^2+^] activity in the final 2.5 s of the optic flow epoch, plotted as a function of the egocentric traveling direction simulated by the optic flow. The translational speed of optic flow increases across the four columns, from left to right. Gray lines: sinusoidal fits. **c**, Same as panel b, but analyzing the right-bridge PFN_d_ activity. **d**, Same as panel b, but analyzing the left-bridge PFN_v_ activity. **e**, Same as panel b, but analyzing the right-bridge PFN_v_ activity. **f**, Same as panel b, but analyzing the hΔB activity in the fan-shaped body. **g**, Same as panel b, but analyzing the PFR activity in the fan-shaped body. **h**, Amplitude of the four sinusoids in panel b to indicate how PFN_d_ responses, overall, scale with optic-flow translation speed. **i-k**, Same as panel h, but for the plots and cell type shown to the left. Note that the amplitudes of the PFN sinusoidal activity patterns are not only scaled by the traveling direction angle (panel b-e), but also by traveling speed (panel h-k). **l**, Mean of the four sinusoids in panel f to indicate how hΔB responses, overall, scale with optic-flow translation speed. **m**, Same as panel l, but for the PFR plots shown to the left. Note that speed tuning of the bolus amplitude in both the hΔBs (panel l) and the PFRs (panel m) is detectable, but weak. Also note that response-scaling with speed in hΔBs and PFRs was not consistent across all traveling directions (panels f and g). **n**, Same as panel b, but analyzing the mean (rather than the amplitude) of the left-bridge PFN_d_ [Ca^2+^] activity patterns. Gray lines: same sinusoidal fits from panel b with a vertical offset and a scale factor that is constant across all four speeds. The fact that our amplitude fits from panel b also fit the mean responses well supports the hypothesis that the heading input and the motion input to PFN cells are integrated multiplicatively (see Methods). **o**-**s**, Same as panel n, but analyzing the cell type indicated on the left side of the figure, for each row. See Methods for how the optic flow speed was calculated.

**Extended Data Figure 9.**
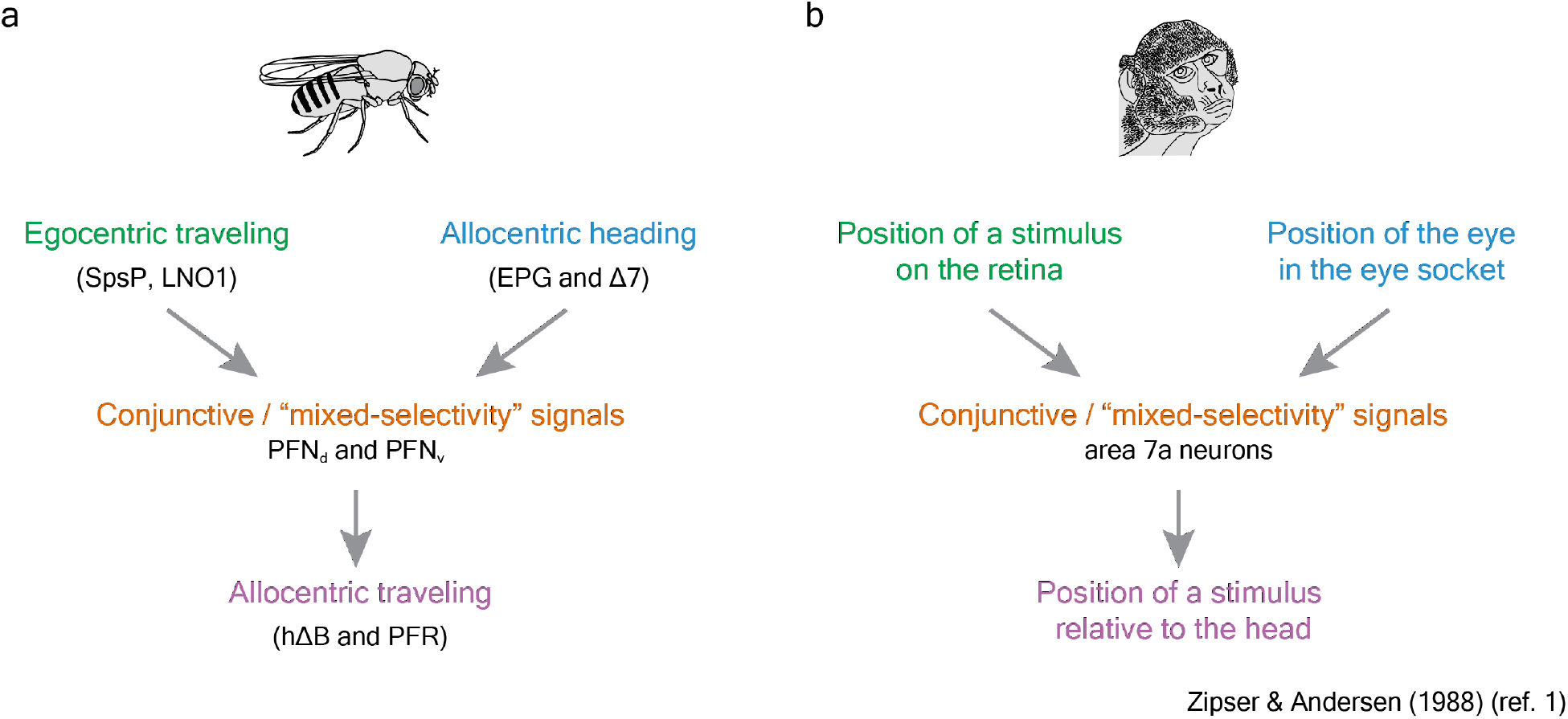
The neural circuit described in this paper implements an egocentric-to-allocentric coordinate transformation. **a**, Schematic of the computation implemented in the fly. Note that traveling-direction signals referenced to the body axis (i.e. optic flow inputs) are converted into traveling-direction signals referenced to cues in the world (i.e. the PFR/hΔB bolus position). **b**, Schematic of a computation hypothesized to take place in the monkey parietal cortex.

**Extended Data Figure 10.**
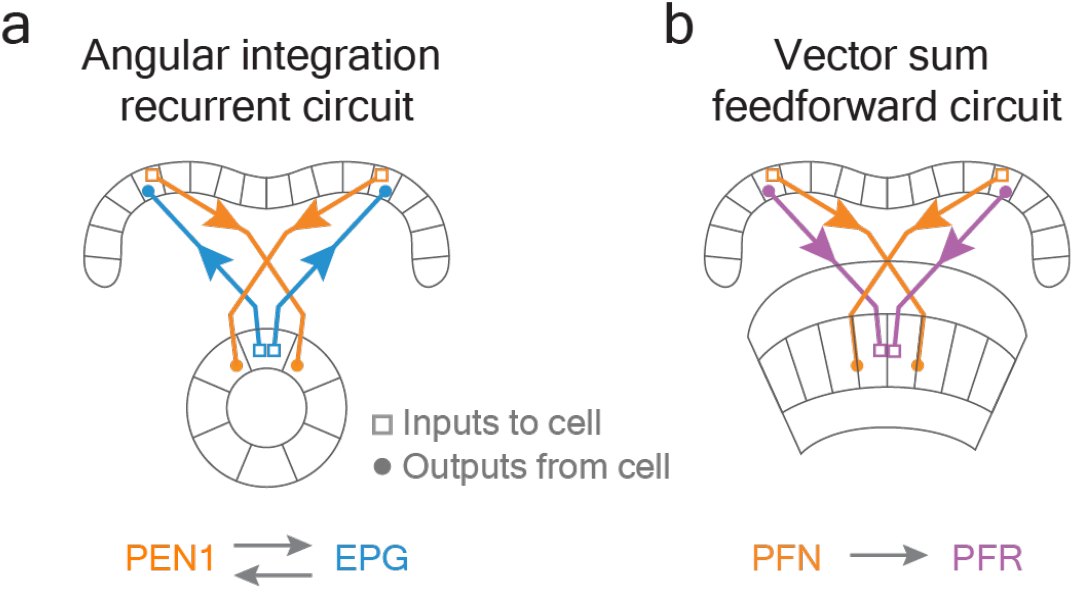
The PFR/PFN system, described in this paper, implements vector addition whereas the anatomically similar EPG/PEN1 system, described previously, implements angular integration. **a**, Left-bridge PEN1 cells project to the ellipsoid body with a 45° clockwise shift relative to EPG cells and right-bridge PEN1 cells project with a 45° counter-clockwise shift. PEN1s are generally thought to electrically signal from the bridge to the ellipsoid body, and EPGs are generally thought to signal in the other direction, from the ellipsoid body to the bridge. This recurrent anatomy is likely what allows the EPG bump to continuously rotate around the ellipsoid body if there is a tonic asymmetry in the left vs. right-bridge PEN1 signal. **b**, Akin to PEN1s, left-bridge PFNs project to the fan-shaped body with a clockwise shift compared to PFRs and right-bridge PFNs project with a counter-clockwise shift. Unlike with the EPG/PEN1 system, both PFNs and PFRs are thought to send electrical signals from the bridge to the fan-shaped body and there is no known columnar cell class that functions as a return path from the fan-shaped body back to the bridge. The PFR/PFN system thus seems to implement a feedforward circuit that performs a vector sum in comparison to the closed-loop EPG/PEN1 system that implements angular integration. In other words, a tonic asymmetry in the left-vs. right-bridge PFNs will yield a tonic offset of the PFR bolus position in the fan-shaped body rather than a continuously rotating bolus, as would be expected with a left-vs. right-bridge signal asymmetry in the EPG/PEN1 system.

**Extended Data Figure 11.**
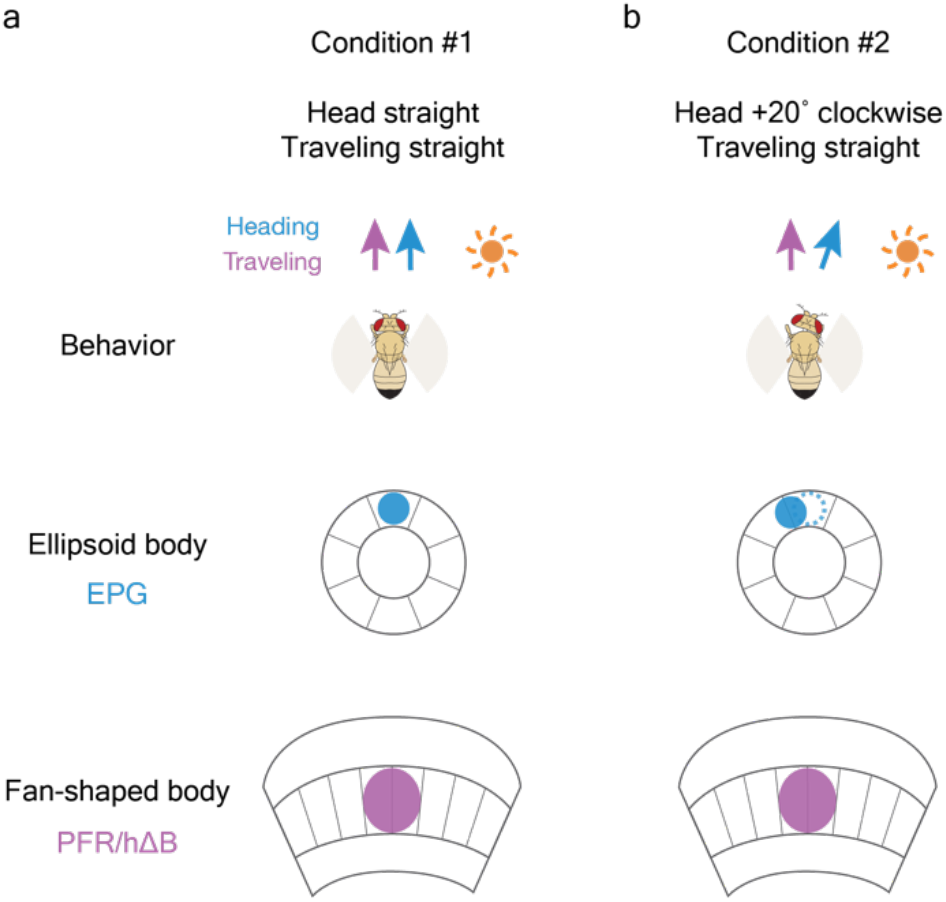
The traveling direction signal computed by optic flow is robust to changes in the angle of the fly’s head. **a**, Fly flying straight with the head aligned to the body axis. EPG and PFR/hΔB signals are aligned in the ellipsoid body and fan-shaped body, respectively. **b**, Fly flying straight forward with the head rotated 20° to the right. The EPG bump––assuming the EPG bump position tracks the fly’s head (rather than body) direction––will rotate 20° counterclockwise. The PFR/hΔB bolus, however, will remain pointing in the same allocentric traveling direction because the net effect of the EPG bump rotating 20° in one direction and the egomotion signal from optic flow (not represented in the diagram) rotating 20° in the opposite direction is that the PFR/hΔB bolus stably indicates the same traveling direction throughout.

